# Targeting the paralog protein for degradation accounts for the opposite roles of NRBP1 and NRBP2 in regulating LINE1 retrotransposition

**DOI:** 10.1101/2024.06.14.598964

**Authors:** Wei Yang, Shaobo Cong, Jennifer Schwarz, Thilo Schulze, Sara Ullrich, Kristina Falkenstein, Denis Ott, Pia Eixmann, Angelica Trentino, Antje Thien, Thierry Heidmann, Ekkehard Schulze, Bettina Warscheid, Ralf Baumeister, Wenjing Qi

## Abstract

Gene duplication generates paralogs undergoing diverse fates during evolution and serves as a potent catalyst of biological complexity. Paralogs frequently share redundant functions and may also exhibit antagonistic activities by competing for common interaction partners. Here we show that the gene paralogs *NRBP1* and *NRBP2* oppositely regulate LINE1 retrotransposition, via influencing integrity of the LINE1 ribonucleoprotein complex. We demonstrate that the opposite roles of NRBP1 and NRBP2 are not results of a competitive mechanism, but rather due to targeting NRBP1 for proteasome-mediated decay by NRBP2, probably through heterodimer formation. Moreover, our phylogenetic analysis shows that the regulatory function of NRBP2 may be acquired later during evolution, suggesting that evolutionary pressure has favored this functional fine-tuning of NRBP1. In summary, our discovery not only identifies NRBP1/2 as novel LINE1 regulators and implicates their involvement in human pathogenesis, but also provides a novel insight into the regulatory details arising from gene duplication.

## Introduction

Paralogs, arising from gene duplication events, play a key role in supplying new genetic material for natural selection in evolution. The functional redundancy shared by paralogs serves as a crucial source of genetic robustness.^1^ Additionally, paralogs may evolve to new functions, either distinct or complementary to those of the original genes.^2^ Despite being the result of gene duplication, the existence of antagonistic functions among paralogs has long been documented.^3-5^ This antagonism is often conceptualized within a competition model, where the protein encoded by a loss-of-function gene copy competes with that from its functional sister copy in binding to common interaction partners.^4^ The persistence and fixation of such an acquired ability, allowing one gene to repress the activity of its sister gene, suggest evolutionary advantages. This phenomenon likely contributes to enhancing the plasticity of complex biological systems, providing an adaptive edge in the dynamic landscape of evolution. Nuclear receptor-binding proteins (NRBPs) are evolutionarily conserved pseudokinases that, despite losing their original function of phosphorylating proteins, can act as allosteric modulators, scaffolds for complex assembly or as competitive inhibitors in signaling pathways.^6^ Although NRBP family members widely exist in metazoan, their molecular functions are not well known. Humans have two NRBP paralogs, *NRBP1* and *NRBP2*. NRBP1 protein has a conserved kinase-like domain that has lost the kinase activity, nuclear export/import signals (NES/NLS), a BC-binding box, a MLF1-binding region and two predicted nuclear receptor binding (NRB) motifs.^7-9^ The BC-binding box has been shown to be responsible for association of NRBP1 with Elongin B/C (ELOB/C).^10^ The MLF1-binding region has been reported to mediate NRBP1 homodimerization which is crucial for assembly of the Elongin B/C containing Cullin-RING E3 ubiquitin ligase complex.^11^ The best studied molecular function of NRBP1 is its participation in proteasome-mediated protein degradation. Here, NRBP1 acts as a substrate recognition receptor in the Elongin B/C containing Cullin-RING E3 ubiquitin ligase complex and has been shown to promote Amyloid β production via accelerating BRI2 and BRI3 degradation in neuronal cells.^11^ In addition, NRBP1 has been found to promote SALL4 degradation and affects various signaling pathways, such as Rac1/Cdc42.^12,13^ In these contexts, NRBP1 exhibits either oncogenic or tumor-suppressive properties to influence tumorigenesis and development.

In contrast to NRBP1, the molecular function of NRBP2 is much less well characterized. NRBP2 was initially recognized for its participation in supporting neural progenitor cell survival.^14^ In addition, NRBP2 is suggested as a tumor suppressor by suppressing key oncogenic signaling pathways, such as AKT and Mammalian Target of Rapamycin (mTOR) pathways.^15,16^ How NRBP2 mechanistically influences signaling transduction or whether it might have a similar molecular function as NRBP1, is currently unknown.

LINE1 (L1) is the only known active autonomous retrotransposon in humans and contributes to about 17% of the human genome.^15,17^ Given that L1 retrotransposition can cause mutations and drive genome instability, its activation is primarily linked to human diseases such as tumors.^15,18^ L1 has also been implicated in several autoimmune diseases by activating the innate immune system.^19,20^ L1 encodes two proteins playing essential roles in successful L1 retrotransposition, an RNA-binding protein open-reading frame 1 (ORF1) and an open-reading frame 2 (ORF2) that possesses endonuclease and reverse transcriptase activities.^21-25^ ORF1 and ORF2 act together to enable mobilization of L1 through a “copy and paste” mechanism. After transcription from the genome and translation of ORF1 and ORF2, L1 mRNA together with ORF1, ORF2 and other RNA-binding proteins assemble into ribonucleoparticles (L1 RNPs) in the cytoplasm.^26-28^ The L1 RNPs subsequently translocate into the nucleus, followed by reverse transcription and integration into new genomic loci.^21,29^ Host cells have developed several defense mechanisms to prevent deleterious L1 mobilization. Most of the L1 DNA copies are inactive due to mutations, rearrangements, or truncations. In addition, DNA methylation and histone modification silence L1 at the transcriptional level.^30^ Furthermore, a variety of ORF1-associated host factors are reported to restrict L1 via different mechanisms.^31,32^ Despite the obvious deleterious consequences of L1 activation, it also has important biological roles under certain circumstances.^33^ L1 activity in the germline is considered to be an important source of genetic diversity.^34,35^ In addition, the L1 transcript has been reported to function as non-coding regulatory RNA and to actively regulate neuronal development and brain function.^36^ Therefore, regulatory mechanisms for both L1 activation and inhibition must have been co-evolved to enable a context- and tissue-specific control of L1.

Here we identify NRBP1 and NRBP2 as novel regulators of L1 retrotransposition. We show that NRBP1 and NRBP2 interact with L1 ORF1 protein but exert opposing effects on L1 mobility. Specifically, NRBP1 activates, whereas NRBP2 restricts L1 retrotransposition by influencing association of L1 mRNA with ORF1 protein. Moreover, we demonstrate that the restrictive role of NRBP2 is achieved by targeting NRBP1 protein for degradation, probably through heterodimer formation between NRBP1 and NRBP2. Furthermore, NRBP2 knock-down results in activation of innate immune response, which is also stimulated upon L1 activation, and NRBP2 expression level displays a negative correlation with Rheumatoid Arthritis (RA) autoimmune disease. Finally, our phylogenetic analysis shows that *NRBP1/2* emerge as gene duplication products of an ancestral *NRBP* in the early vertebrate lineage. *NRBP1* probably maintains the functions of its ancestral *NRBP*, while *NRBP2* may have obtained new regulatory roles during its evolution. In summary, this study not only uncovers the opposite roles of NRBP1 and NRBP2 in regulating L1 retrotransposition, but also provides a mechanistic insight about how one protein inhibits its paralogous protein via heterodimerization-dependent protein degradation.

## Results

### NRBP1 and NRBP2 interact with L1-encoded ORF1

As catalytically inactive enzymes, pseudokinases can function as scaffold proteins to mediate protein interactions. Therefore, identifying binding partners of NRBP1 and/or NRBP2 (NRBP1/2) would shed light on the protein complexes wherein NRBP1/2 carry out their functions. Therefore, we immunoprecipitated endogenous NRBP1 and NRBP2 in MCF-7 cells by using an antibody recognizing both pseudokinases (**Figure S1A**) and identified co-immunoprecipitated proteins via Liquid Chromatography–Mass Spectrometry (LC–MS). In total, we identified 107 proteins as associated with NRBP1/2 (enrichment *vs.* mock control ≥ 5 fold, p-value < 0.05), including the previously known NRBP1 interactors: ELOB (TCEB2), ELOC (TCEB1), TSC22D1 and TSC22D2 (**Figure 1A**; **Table S1**). In addition, we noticed that ORF1, which is encoded by the L1 retrotransposon, was co-immunoprecipitated with NRBP1/2. Moreover, multiple proteins that are known to regulate L1 retrotransposition or associate with ORF1 were also identified as NRBP1/2 interactors (**Table S1**). We next confirmed the interaction of endogenous NRBP1/2 with ORF1 via Co-IP using NRBP1/2 antibody (**Figure 1B**). Transfection of MCF-7 cells with Myc-Flag-NRBP1 or Myc-Flag-NRBP2 allowed us to pull down endogenous ORF1 with anti-Flag antibody (**Figure 1C**). Flag-tagged ORF1 also enabled pull-down of both NRBP1-Myc and NRBP2-Myc (**Figure 1D**). These results together suggest that both NRBP1 and NRBP2 interact with ORF1. Furthermore, we confirmed the association of both NRBP1 and NRBP2 with some known ORF1 interactors that are also identified in our MS analysis, including UPF1, MOV10, G3BP1 and YBX1 (**Figure S1B**), suggesting that NRBP1 and NRBP2 might be novel components of the previously described L1 RNP complexes that affect L1 mobility.

**Figure 1.**
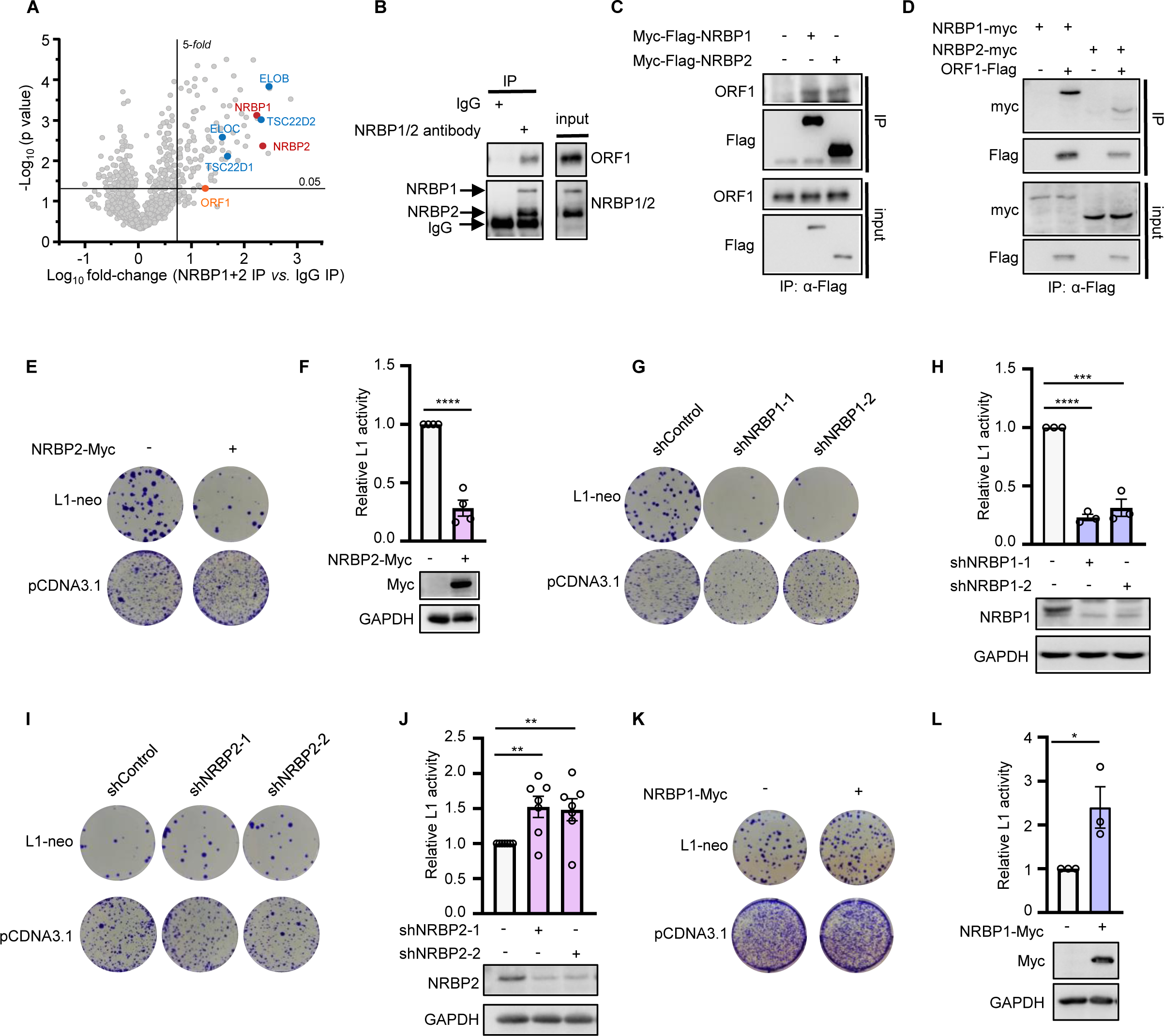
NRBP1 and NRBP2 interact with L1-encoded ORF1 and contrarily regulate L1 retrotransposition. **(A)** Scatter plots showing proteins significantly enriched (fold change ≥ 5, *p*-value < 0.05) in NRBP1/2 IP *vs.* IgG in MCF-7 cells. NRBP1 and NRBP2 are shown in red. Previously known NRBP1 interactors are labelled in blue. ORF1 is marked in orange. **(B)** Endogenous ORF1 is co-immunoprecipitated with NRBP1 and/or NRBP2. An antibody recognizing both NRBP1 and NRBP2 was used to pull down endogenous NRBP1 and NRBP2 in MCF-7 cells. Co-precipitated ORF1 was detected by using ORF1 antibody. **(C)** Endogenous ORF1 is co-immunoprecipitated with both NRBP1 and NRBP2. MCF-7 cells were transfected with either Myc-Flag-tagged NRBP1 or Myc-Flag-tagged NRBP2. The cell lysates were incubated with Flag antibody and the immunoprecipitated proteins were detected by Western blot with the indicated antibodies. **(D)** NRBP1 and NRBP2 are co-immunoprecipitated with ORF1. Flag antibody was used to pull down ORF1-Flag, and Myc antibody was used to detect co-immunoprecipitated Myc-NRBP1 and Myc-NRBP2 in HeLa cells. **(E-L)** NRBP1 activates and NRBP2 inhibits L1 retrotransposition. One representative L1 retrotransposition activity is shown in **(E)**, **(G)**, **(I)** and **(K)**. Quantitation of L1 retrotransposition activity is shown in the top panels of **(F)**, **(H)**, **(J)** and **(L)**. Western blots indicating successful overexpression or knockdown are shown in the bottom panels of **(F)**, **(H)**, **(J)** and **(L)**. N = 4 biological replicates for **(E)** and **(F)**. N = 3 biological replicates for **(G)** and **(H)**. N = 7 biological replicates for **(I)** and **(J)**. N = 3 biological replicates for **(K)** and **(L)**. See also Figure S1 and Table S1.

### NRBP1 and NRBP2 regulate L1 retrotransposition in opposite ways

The formation of the cytosolic L1 RNP complex, initiated by the binding of L1 mRNA to ORF1, is a crucial prerequisite for L1 retrotransposition. Many regulators of L1 retrotransposition exert their regulatory roles through interacting with ORF1.^31,32,37,38^ We asked whether NRBP1 or NRBP2 might influence L1 retrotransposition. Successful retrotransposition results in cellular resistance to G418 neomycin in cells expressing a L1-neo reporter.^39^ We co-transfected HeLa cells with the L1-neo reporter and NRBP1-myc or NRBP2-myc expressing constructs followed by G418 selection. We also transfected L1-neo reporter into the NRBP1 or NRBP2 shRNA knockdown HeLa cell lines. The results of the colony assay showed that either NRBP2 overexpression or NRBP1 knockdown strongly reduced L1 retrotransposition (**Figure 1E-H**), while NRBP2 knockdown or NRBP1 overexpression increased L1 activity (**Figure 1I-L**). These observations together suggest NRBP1 as a positive and NRBP2 as a negative regulator of L1 retrotransposon.

### NRBP1 and NRBP2 contrarily regulate ORF1 and L1 mRNA association

Altering subcellular localization of L1 RNP complexes can affect L1 retrotransposition.^40,41^ Since both NRBP1 and NRBP2 interact with ORF1 protein and regulate L1 activity, we asked whether they could affect subcellular localization of ORF1. As MCF-7 cells have a higher endogenous ORF1 expression level compared to HeLa cells (**Figure S2A**), we examined localization of endogenous ORF1 in MCF-7 cells. ORF1 in MCF-7 cells exhibited a predominantly diffuse distribution, accompanied with enrichment in a few small punctate structures in the cytoplasm (**Figure 2A**). The number of ORF1 containing puncta increased upon knockdown of NRBP1 (**Figure 2A**). Sequestration of ORF1 and L1 mRNA into stress granules (SGs) was shown before to inhibit L1 retrotransposition.^40^ As the ORF1 containing puncta upon NRBP1 knockdown did not show an enrichment of the SG marker G3BP1 (**Figure 2A**), they are probably not SGs.

**Figure 2.**
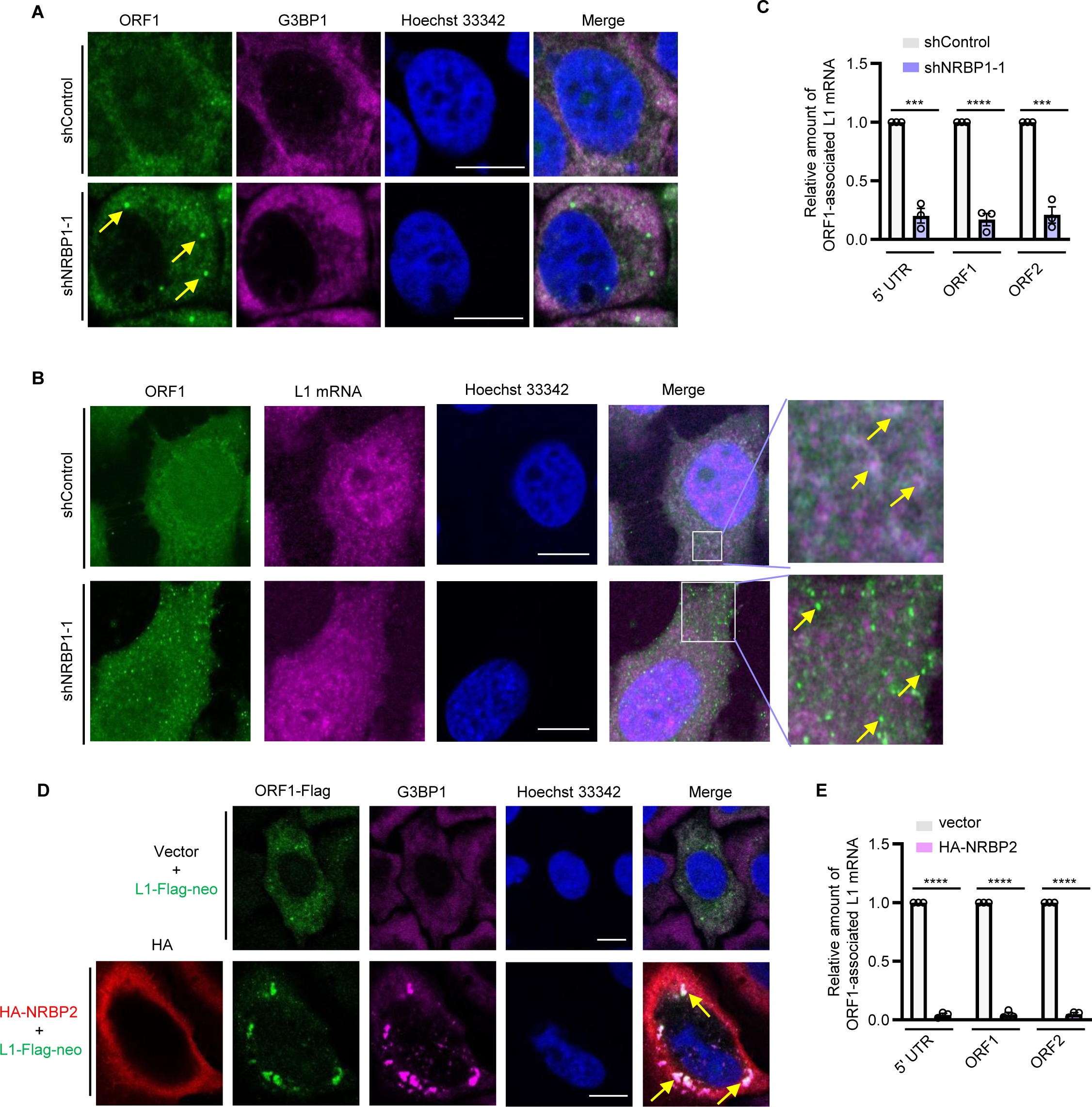
NRBP1 and NRBP2 contrarily regulate ORF1 and L1 mRNA association. **(A)** NRBP1 knockdown results in an enrichment of ORF1 in cytoplasmic foci in MCF-7 cells. The yellow arrows point to the ORF1 foci. Shown are immunofluorescence staining of endogenous ORF1 and G3BP1 proteins with their respective antibodies. Scale bar 10 µm. **(B)** NRBP1 knockdown impairs colocalization of ORF1 and L1 mRNA. Shown are immunofluorescence staining of endogenous ORF1 protein with ORF1 antibody (green) and L1 mRNAs detected via smFISH with L1 probes (violet). Nucleus was stained with Hoechst 33342 (blue). The arrows in the top panel point to co-localized ORF1 and L1 mRNA, the arrows in the bottom panel point to the ORF1-enriched puncta without an enrichment of L1 mRNA. Scale bar 10 µm. **(C)** ORF1 RIP-qPCR to quantitate ORF1-associated L1 mRNA. Three primer pairs targeting to L1 5′ UTR, ORF1 and ORF2 were used for qPCR. N = 3 biological replicates. **(D)** HA-NRBP2 overexpression results in an enrichment of ORF1-Flag in cytoplasmic puncta in HeLa cells. The yellow arrows point to colocalized ORF1 and G3BP1 foci. Scale bar 10 µm. **(E)** NRBP2 overexpression prevents the interaction between ORF1 protein and L1 mRNA. RIP-qPCR was carried out to quantitate Flag-tagged ORF1 co-immunoprecipitated L1 mRNA levels with indicated primer pairs. N = 3 biological replicates. See also Figure S2.

Association of L1 mRNA with ORF1 protein to form the L1 RNP complex is a prerequisite of its retrotransposition life cycle. We wondered whether NRBP1 could affect L1 RNP complex assembly. We then used single molecule fluorescence in situ hybridization (smFISH) to detect endogenous L1 mRNA in MCF-7 cells and examined the influence of NRBP1 on subcellular localization of L1 mRNA and ORF1 protein. In control cells, we observed a partial colocalization of ORF1 and L1 mRNA in cytoplasm, while NRBP1 knockdown led to a strongly decreased colocalization of the protein and mRNA (**Figure 2B**). Especially in the ORF1 puncta, an enrichment of L1 mRNA was not obvious. This observation raised the question whether NRBP1 knockdown results in a disassociation of ORF1 protein and L1 mRNA. To answer this question, we used ORF1 antibody to pull down endogenous ORF1 and quantified the co-precipitated L1 mRNA by RT-qPCR (RIP-qPCR). Three pairs of primers targeting different parts of L1 mRNA were used (5′ UTR, ORF1-coding region, ORF2-coding region). We found that NRBP1 knockdown strongly reduced the amount of L1 mRNA co-immunoprecipitated with ORF1 (**Figure 2C and S2B**), suggesting an impaired binding of ORF1 protein to L1 mRNA in the absence of NRBP1.

As the inhibitory role of NRBP2 on L1 is more prominent upon NRBP2 overexpression than knockdown, we checked influence of NRBP2 overexpression on L1 ORF1 localization. We transfected HeLa cells with HA-tagged NRBP2 and L1-Flag-neo plasmid. Similar to endogenous ORF1 in MCF-7 cells, Flag-tagged ORF1 in HeLa cells was mostly diffusely localized in the cytoplasm, with a few enrichments of cytoplasmic puncta (**Figure 2D**). HA-NRBP2 transfection resulted in translocation of ORF1 into large irregular puncta, without an obvious enrichment of HA-NRBP2 itself in these cellular structures (**Figure 2D**). These puncta were larger than those formed in control or NRBP1 knock-down cells and resembled the morphology of SGs (**Figure 2A, B, D**). Indeed, SG marker G3BP1 was also enriched in these L1 ORF1 puncta upon HA-NRBP2 overexpression (**Figure 2D**). G3BP1 plays an essential role in SG assembly and G3BP1 knockdown has been shown to lead to the loss of large ORF1 foci, which in turn activates L1 retrotransposition.^40^ Since G3BP1 was an NRBP1/2 interactor according to our MS and Co-IP results (**Figure S1B, Table S1**), we wondered whether NRBP2 might facilitate SG assembly, leading to the subsequent sequestration of ORF1 into SGs and, thus, inactivation of L1. We generated G3BP1 knockout cell lines to check whether formation of the L1 ORF1 puncta upon NRBP2 overexpression was dependent on G3BP1. NRBP2 still promoted translocation of ORF1 into puncta in the absence of G3BP1 (**Figure S2C**) and inhibited L1 retrotransposition (**Figure S2D, E**). These data implicate that ORF1 protein upon NRBP2 overexpression shuttles into certain cellular puncta whose biogenesis is independent of G3BP1, and NRBP2 inhibits L1 retrotransposition independent of the G3BP1-mediated SG pathway. To determine whether NRBP2 overexpression also results in ORF1 and L1 mRNA disassociation, we performed ORF1 RIP-qPCR. NRBP2 overexpression significantly reduced the ability of Flag-tagged ORF1 to pull-down L1 mRNA, indicating a strongly reduced affinity of ORF1 to L1 mRNA (**Figure 2E and S2F**).

In summary, our results suggest that NRBP1 may facilitate and NRBP2 may inhibit ORF1 interaction with L1 mRNA, thereby exerting opposite roles in L1 retrotransposition.

### NRBP2 negatively regulates NRBP1 to inhibit L1 retrotransposition

Opposite functions among protein encoded by paralogs have been described, frequently due to competition for interaction with common binding partners.^4^ As both NRBP1 and NRBP2 interact with ORF1, we asked whether presence of one NRBP might abrogate interaction between ORF1 and the other NRBP. We found that neither NRBP2 nor NRBP1 exerted a significant influence on the affinity of the other NRBP with ORF1 (**Figure S3A**), arguing against such a model of competition between NRBPs in ORF1 binding. Next, we asked whether one of these NRBPs may regulate L1 by inhibiting their respective counterpart. In such a scenario, the influence of the upstream inhibitor would be nullified in the absence of the downstream factor. We found that NRBP2 failed to inhibit L1 activity upon NRBP1 knockdown (**Figure 3A and S3B**), indicating that NRBP2 might inactivate L1 via inhibiting NRBP1. In addition, we noticed that NRBP2 overexpression reduced the protein level of endogenous NRBP1 in HeLa cells (**Figure 3B**) and NRBP2 knockdown resulted in an elevated NRBP1 protein level without affecting NRBP1 mRNA level (**Figure 3C, D and S3C**). Notably, NRBP1 knockdown reversely increased both NRBP2 mRNA and protein levels (**Figure 3C and S3D, E**). These observations together implicate that NRBP1 and NRBP2 inhibit expression level of their respective paralogs, but via distinct mechanisms: while NRBP1 probably affects transcription or mRNA stability of NRBP2, NRBP2 negatively influences NRBP1 activity at translational or post-translational level. A frequently observed mechanism of post-translational inhibition involves redirecting a protein to a different subcellular location, so we asked whether NRBP2 can alter NRBP1 subcellular distribution. Consistent with the previous reports,^42^ single transfected HA-NRBP1 in HeLa cells displayed an even distribution of the protein in cytoplasm with a perinuclear reticular staining pattern which might hint at its localization on the ER membrane (**Figure 3E**). In contrast, overexpressed NRBP2-Myc alone displayed a diffused localization in the cytoplasm without such a reticular staining pattern. In cells co-transfected with both HA-NRBP1 and NRBP2-Myc, HA-NRBP1 became more enriched in the peripheral region, while NRBP2-Myc did not show obvious alteration of its subcellular localization (**Figure 3E**). In summary, our results suggest that NRBP2 not only reduces protein level but also influences subcellular localization of NRBP1.

**Figure 3.**
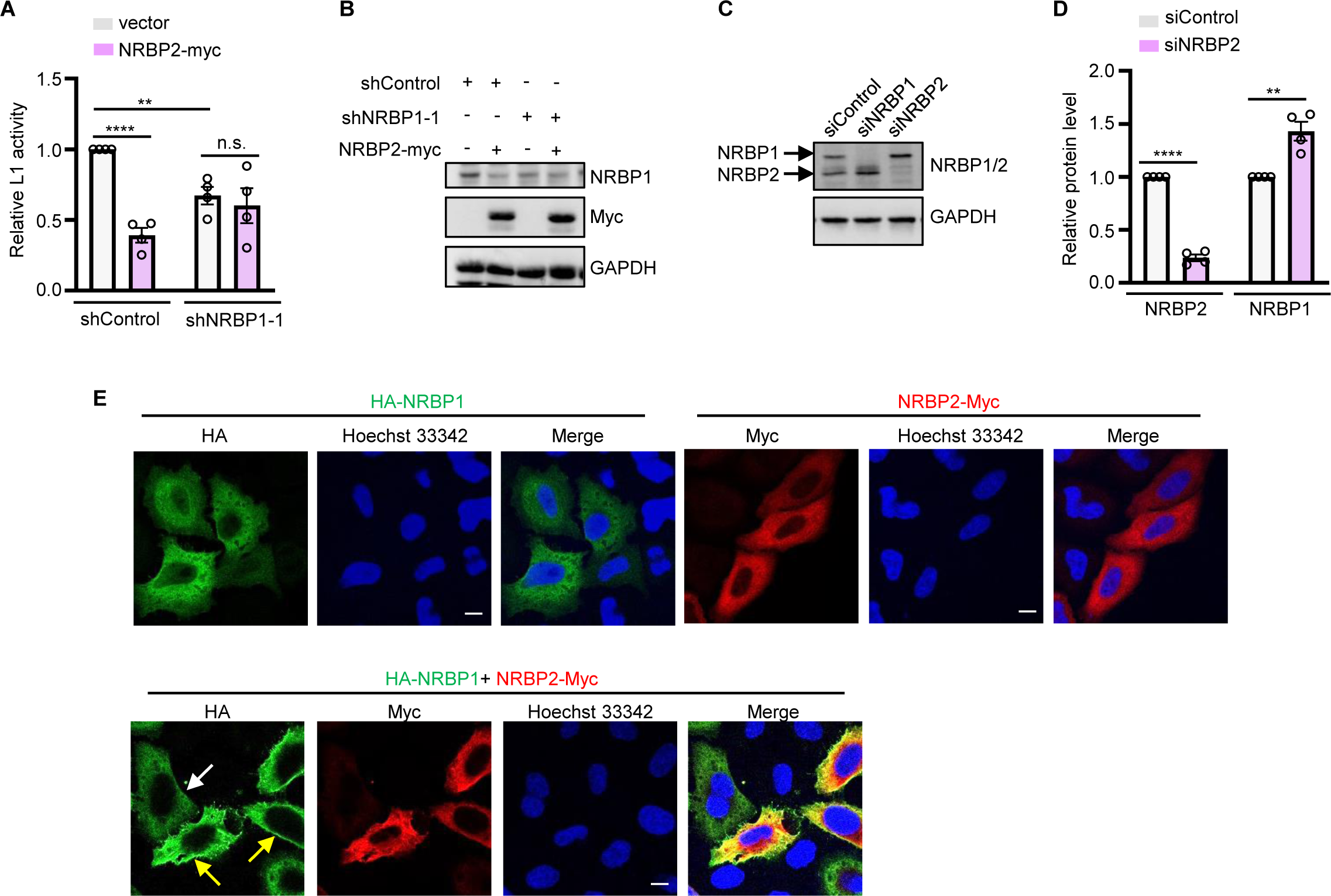
NRBP2 inactivates L1 retrotransposition via NRBP1 inhibition. **(A)** NRBP1 knockdown abolishes the inhibitory effect of NRBP2 on L1 in HeLa cells. Shown is quantitation of L1 retrotransposition activity. N = 4 biological replicates. **(B)** Overexpression of NRBP2-Myc reduces endogenous NRBP1 protein level. Shown is one representative Western blot using the samples from **(A)** to detect endogenous NRBP1 and transfected NRBP2-Myc. **(C and D)** NRBP1 and NRBP2 negatively affect the protein levels of each other in HeLa cells. Endogenous NRBP1 and NRBP2 were detected using NRBP1/2 antibody. Shown is one representative Western blot result **(C)**. Quantification of the protein levels was shown in **(D)**. N = 4 biological replicates. **(E)** Overexpressing NRBP2-Myc results in an enrichment of HA-NRBP1 to the peripheral region of HeLa cells. Yellow arrows point to cells co-expressing HA-NRBP1 and NRBP2-Myc. White arrow indicates a cell only expressing NRBP1. HA-NRBP1 and NRBP2-Myc were detected using HA and Myc antibodies. Scale bar 10 µm. See also Figure S3.

### The C-terminal halves of both NRBP2 and NRBP1 negatively regulate L1 retrotransposition

Differences in the amino acid sequences, and thus structure, of NRBP2 should account for its inhibitory activity on NRBP1. We therefore compared these two closely related proteins. Alignment of the protein sequences revealed that NRBP1 and NRBP2 share 55.7% amino acid identity. In addition, most of the predicted structures of NRBP1 are conserved in NRBP2. We noticed that NRBP1 has a longer N-terminus (1-43 aa) than NRBP2. This region is a ‘simple sequence’, containing more than 30% serine. A part of it is recognized by the Segmasker algorithm, which detects low complexity sequence (**Figure S4 and S5A**).^43^ Additionally, this region (1-43 aa) is unstructured according to Alphafold, resembling the feature of a low complexity region (LCR) (**Figure S5B**).^44,45^ We asked whether this LCR might account for the functional difference between NRBP1 and NRBP2. To answer this question, we generated an NRBP1 mutant without the LCR (ΔLCR-NRBP1) and a chimeric LCR-NRBP2 mutant by fusing the LCR of NRBP1 to the N-terminus of NRBP2 to test their impacts on L1 retrotransposition (**Figure S5C**). We found that these mutants still behaved similarly as their wild-type counterparts (**Figure S5C-E**), suggesting that the N-terminal LCR does not play a critical role in discriminating NRBP1 from NRBP2.

We reasoned that elucidating critical regions of NRBP1/2 in L1 regulation would be indicative to answer how NRBP2 might inhibit NRBP1. We generated different NRBP2 mutant proteins via eliminating certain domains or motifs and tested their regulatory roles on L1 (**Figure 4A**). While NRBP2 mutants lacking the proposed nuclear export signal (NES) or nuclear localization signal (NLS) could still inhibit L1, eliminating the proposed NRB motif (dNRB) or the dimerization region (dDimer) abolished the inhibitory effect of NRBP2. A variant of NRBP2 lacking the Elongin BC-binding motif (dBC) displayed a partial loss of function (**Figure 4B and S5F**). As BC-binding, NRB and dimerization motifs are all localized in the C-terminal half of the NRBP2 protein, we wondered whether this region might exert the inhibitory role. We next tested whether overexpression of N-terminal or C-terminal halves of NRBP2 (NRBP2-N and NRBP2-C, illustrated in **Figure 4A**) alone could affect L1 activity. NRBP2-C conferred an even stronger suppressive effect on L1 retrotransposition than wild-type NRBP2, despite its significantly lower expression level. In contrast, NRBP2-N failed to inhibit L1 (**Figure 4C and S5G**). These data together suggest that C-terminal half of NRBP2 is necessary and sufficient to inhibit L1. We next asked whether loss of inhibition in NRBP2-N is caused by its loss of ORF1 binding. We found both NRBP2-N and NRBP2-C could be co-immunoprecipitated with ORF1, implicating that NRBP2 might interact with ORF1 through multiple interfaces, and interaction alone is not sufficient for NRBP2-N to restrict L1 retrotransposition (**Figure S5H**).

**Figure 4.**
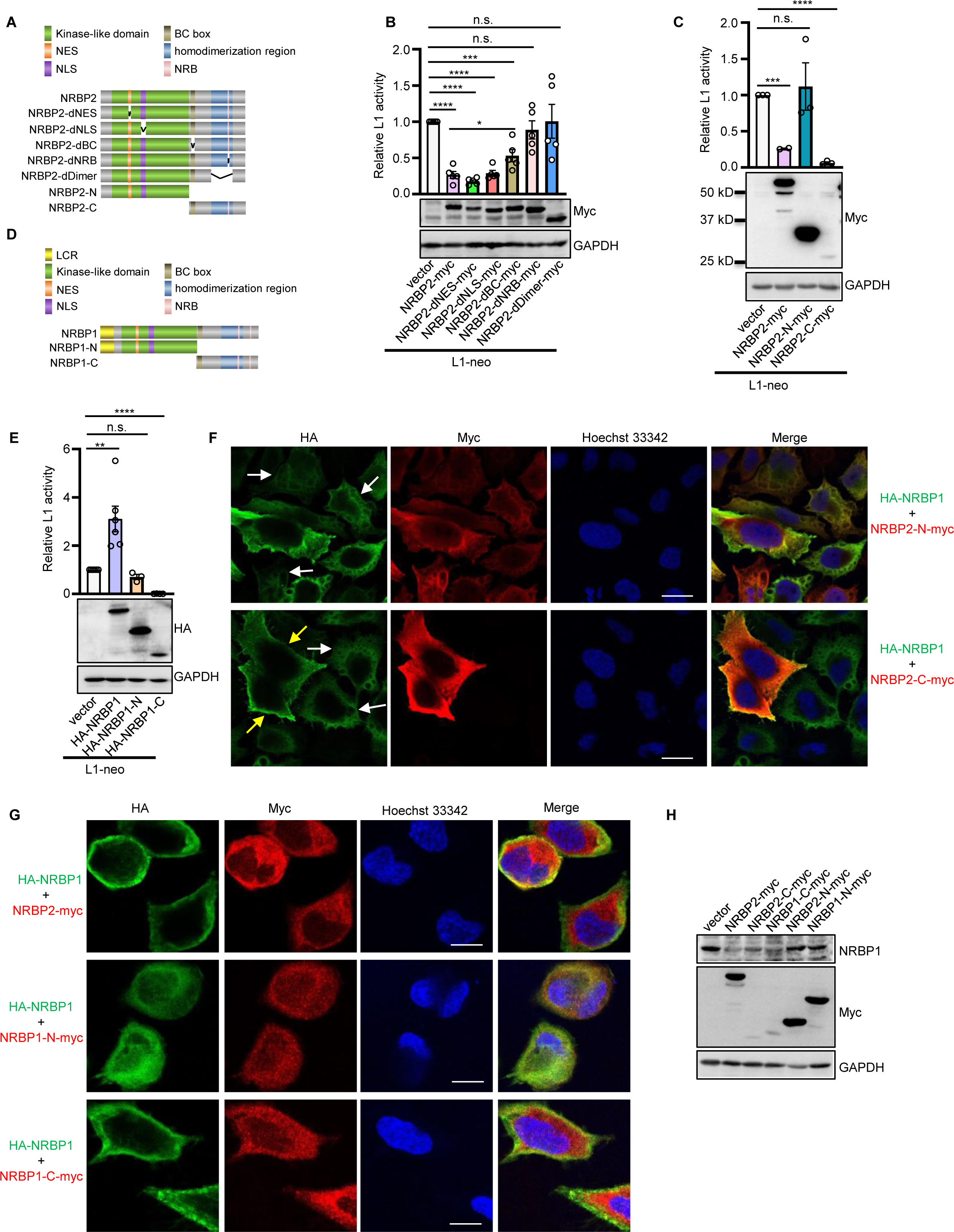
The C-terminal halves of NRBP2 and NRBP1 inhibit L1 retrotransposition and drive full-length NRBP1 to the peripheral region of cells. **(A)** Schematic diagrams of NRBP2 and the NRBP2 mutants used in this study. NES: nuclear export signal; NLS: nuclear localization signal; BC box: Elongin BC-binding motif; NRB: nuclear receptor-binding motif. **(B)** The NRB and dimerization region of NRBP2 are required for NRBP2 to inhibit L1 retrotransposition. Quantification of L1 activity is shown in the top panel. N = 5 biological replicates. One representative Western blot showing expression of the respective NRBP2 mutant variants is shown in the bottom panel. **(C)** The C-terminal half of NRBP2 is necessary and sufficient to inhibit L1. Shown is quantification of L1 activity upon expression of the respective NRBP2 variants. N = 3 biological replicates. **(D)** Schematic diagram of NRBP1 and the NRBP1 halves used in this study. LCR: low complexity region; NES: nuclear export signal; NLS: nuclear localization signal; BC box: Elongin BC-binding motif; NRB: nuclear receptor-binding motif. **(E)** The C-terminal half of NRBP1 functions oppositely to the full-length NRBP1 and inhibits L1. Shown is quantification of L1 activity upon expression of the respective NRBP1 variants. For NRBP1, N = 5 biological replicates. For NRBP1-N and NRBP1-C, N = 3 biological replicates. **(F)** The C-terminal half of NRBP2 promotes enrichment of NRBP1 to the peripheral region of HeLa cells. White arrows point to NRBP1 without NRBP2 transfection. Yellow arrows indicate cells with enrichment of NRBP1 to the peripheral region upon NRBP2 overexpression. Scale bar 10 µm. **(G)** The C-terminal half of NRBP1 promotes enrichment of full-length NRBP1 to the peripheral region of HeLa cells. Scale bar 10 µm. **(H)** Overexpressing NRBP2 and C-terminal half of NRBP1 or NRBP2 reduces endogenous NRBP1 protein level in HeLa cells. Shown is one representative Western blot to detect the respective NRBP1 or NRBP2 variants and endogenous NRBP1. N = 3 biological replicates. See also Figure S4, S5 and S6.

Since NRBP1 and NRBP2 have opposing activities in L1 regulation, we wondered whether the C-terminal halves of these two proteins are sufficient for these divergent functions. We constructed NRBP1-N and NRBP1-C halves (**Figure 4D**) and found that NRBP1-N, as expected, did not show any effect on L1 retrotransposition. To our surprise, NRBP1-C strongly repressed L1 retrotransposition, behaving similarly as NRBP2-C but oppositely to the full-length NRBP1 (**Figure 4E and S5I**). These results indicate that the presence of the N-terminal half switches the functionality of the C-terminal half of NRBP1.

Next, we tested whether NRBP2-C or NRBP1-C, which both inhibit L1 activity, would behave similarly as full-length NRBP2 to regulate protein level and subcellular distribution of full-length NRBP1. Transfection of NRBP2-C-Myc or NRBP1-C-Myc resulted in an enrichment of full-length HA-NRBP1 in the cell periphery and a decrease of NRBP1 protein level (**Figure 4F-H and S6**), similar as full-length NRBP2. In contrast, overexpressing of NRBP2-N-Myc or NRBP1-N-Myc, which alone did not affect L1 activity, altered neither subcellular distribution nor protein level of NRBP1 (**Figure 4F-H and S6**). All together, these data suggest that the isolated C-terminal halves of NRBP1 and NRBP2 exert a similar function as NRBP2 to inhibit full-length NRBP1, thereby preventing L1 retrotransposition.

### NRBP2 inhibits NRBP1 protein expression via proteasome-mediated degradation

Our results so far demonstrate that NRBP2 not only negatively affects protein level of NRBP1 without affecting its mRNA level, but also promotes a distribution of NRBP1 to the cell periphery. There are two scenarios of how NRBP2 could accomplish this NRBP1 regulation. In the first model, NRBP1 is redirected to the periphery by NRBP2, where it is functionally inactive. In the second model, degradation of NRBP1 in the nuclear periphery is stimulated by NRBP2, and the peripheral remaining NRBP1 is insufficient to execute its L1 activating function. To test these two possibilities, we asked whether triggering degradation of NRBP1 in the perinuclear region by NRBP2 could account for the change in both protein level and subcellular localization of NRBP1. Overexpression of NRBP2 failed to reduce NRBP1 protein level in the presence of protease inhibitor MG132 (**Figure 5A and S7A**). Similarly, both NRBP1-C and NRBP2-C reduced NRBP1 protein level and this could be blocked with MG132 (**Figure 5A and S7A**). In contrast, NRBP1-N and NRBP2-N, which did not affect L1 retrotransposition, had no effect on NRBP1 protein level (**Figure 5A and S7A**). Consistently, we found that when NRBP1 was co-expressed with either NRBP2-C or NRBP1-C, it showed a preference of localization in the peripheral region of cells and MG132 partially restored its localization in the perinuclear region (**Figure 5B**). These observations together indicate that the predominant peripheral localization of NRBP1, triggered by NRBP2 or the C-terminal halves of both NRBPs, is probably a consequence of NRBP1 degradation in the perinuclear region of the cell.

**Figure 5.**
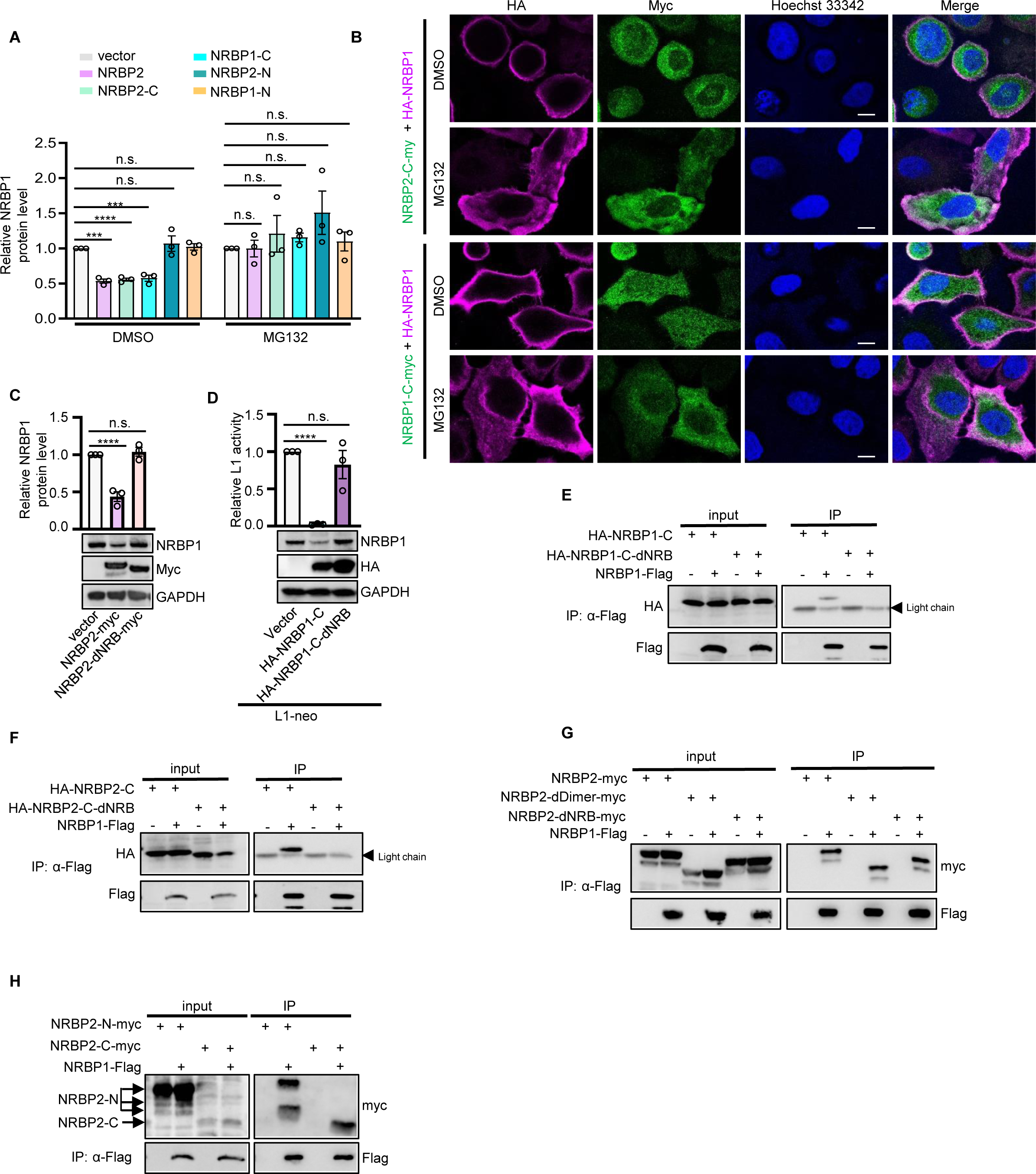
NRBP2 promotes proteasome-mediated degradation of NRBP1 protein. **(A)** Proteasome inhibitor MG132 abolishes capability of NRBP2 and the C-terminal halves of NRBP1 and NRBP2 to reduce protein level of NRBP1. Shown is quantification of relative protein levels of NRBP1 in HeLa cells. GAPDH was used for normalization. N = 3 biological replicates. **(B)** Proteasome inhibitor MG132 prevents enrichment of NRBP1 to the peripheral region of HeLa cells upon overexpressing the C-terminal half of NRBP1 or NRBP2. Shown are representative confocal images of immunofluorescence stained cells. Scale bar 10 µm. **(C)** NRB motif in NRBP2 is essential to reduce protein level of NRBP1 in HeLa cells. The upper panel shows the quantification of the relative NRBP1 protein level normalized by GAPDH. Bottom panel is one representative Western blot used for quantification. N = 3 biological replicates. **(D)** NRB motifs in C-terminal half of NRBP1 are essential to inhibit L1 and to reduce the protein level of NRBP1. Quantification of L1 activity in HeLa cells is shown in the top panel. N = 3 biological replicates. One representative Western blot showing expression of the respective proteins is shown in the bottom panel. **(E)** The C-terminal half of NRBP1 interacts with the full-length NRBP1 in an NRB-dependent manner. **(F)** The C-terminal half of NRBP2 interacts with the full-length NRBP1 in an NRB-dependent manner. **(G)** Lack of NRB and dimerization region does not prevent interaction between NRBP2 and NRBP1. **(H)** Both the N- and C-terminal halves of NRBP2 interact with NRBP1. For **E-H**, HEK293T cells were transfected with the indicated plasmids. Flag antibody was used to pull down Flag-NRBP1. The co-precipitated proteins were detected by using HA or Myc antibody. See also Figure S6 and S7.

NRBP1 is known to act as the substrate recognition factor in the Elongin B/C E3 ubiquitin ligase complex to facilitate protein degradation,^11^ while a role of NRBP2 in protein turn-over has not been described before. An appealing possibility would be that NRBP2 could trigger degradation of NRBP1 via an Elongin B/C dependent way. However, knockdown of Elongin B or Elongin C did not prevent NRBP1 degradation upon overexpression of NRBP2 or NRBP1-C, NRBP2-C (**Figure S7B, C**), suggesting that they probably target NRBP1 for decay independently of the Elongin B/C E3 ubiquitin ligase complex.

### Targeting NRBP1 to degradation requires its interaction with the C-terminal half of either NRBP1 or NRBP2

We further explored how NRBP2 could promote protein degradation of NRBP1. As the functional C-terminal halves of both NRBP1 and NRBP2 contain the dimerization regions, we wondered whether NRBP2 and NRBP1-C might interact with the full-length NRBP1 to promote its degradation. Given the important role of the NRB motif of NRBP2 in regulating L1, we also tested the respective mutants with NRB deletions. Consistent with its inability to regulate L1, NRBP2-dNRB failed to reduce NRBP1 protein level (**Figure 5C**). Similarly, NRBP1-C without the two NRB motifs was incapable of decreasing NRBP1 protein level and L1 activity (**Figure 5D and S7D**), confirming the significance of the NRB motifs for both NRBP2 and NRBP1-C in regulating NRBP1 protein level and L1 retrotransposition. Moreover, our Co-IP result showed that both NRBP1-C and NRBP2-C were co-immunoprecipitated with full-length NRBP1 and this required their NRB motifs (**Figure 5E, F**). These observations together suggest that the NRB motifs are essential for the C-terminal halves of both NRBPs to bind to full-length NRBP1 and to promote NRBP1 degradation. In contrast to our expectation, NRBP1 interacted with both wild-type NRBP2 and the NRBP2 mutants lacking the dimerization region or the NRB motif (**Figure 5G**), implicating an involvement of additional motifs or domains in NRBP2 to NRBP1 interaction. Indeed, the N-terminal half of NRBP2 was also immunoprecipitated with NRBP1 (**Figure 5H**). Taken together, our data demonstrated that both N- and C-terminal halves of NRBP2 interact with NRBP1 and the most likely consequence is that they form heterodimers. However, only the NRB-dependent interaction at the C-termini is necessary and sufficient to trigger NRBP1 degradation.

### Down-regulation of NRBP2 activates inflammatory and type I Interferon pathways

L1 activation provokes innate immune response and stimulates type I interferon (IFN) and inflammatory genes.^46,47^ Several negative regulators of L1 have been found to repress innate immune response via L1 inhibition.^38,48,49^ We asked whether NRBP2 could also have a similar impact on genes involved in innate immunity. We performed mRNA-seq and checked alteration of gene expression in response to NRBP2 knockdown. The mRNA-seq in HeLa cells revealed that NRBP2 knockdown led to increased expression of 427 genes and reduced expression of 335 genes (fold change ≥ 1.5, FDR < 0.05, **Figure 6A**; **Table S2**). Gene Ontology (GO) term analysis showed that genes activated upon NRBP2 knockdown were enriched in innate immune response (**Table S2**). In contrast, NRBP1 knockdown had a less prominent impact on gene expression, resulting in 99 upregulated genes and 54 downregulated genes, without significantly affecting the expression of immune-related genes (**Figure 6B**; **Table S2**). To learn whether the elevated expression of the innate immunity genes upon NRBP2 knockdown is linked to L1 activation, we compared our mRNA-seq result with that from a previous study which investigated transcriptional alteration caused by overexpressing L1.^49^ We found that 26% of the genes upregulated by NRBP2 knockdown were also activated upon L1 overexpression (**Figure 6C**) and these overlapping genes were mostly enriched in immune response (**Figure 6D**; **Table S2**). We confirmed activation of some of these immune responsive genes upon NRBP2 knockdown by qRT-PCR (**Figure 6E**). These data together indicate that activation of innate immune response upon NRBP2 knockdown might be caused by L1.

**Figure 6.**
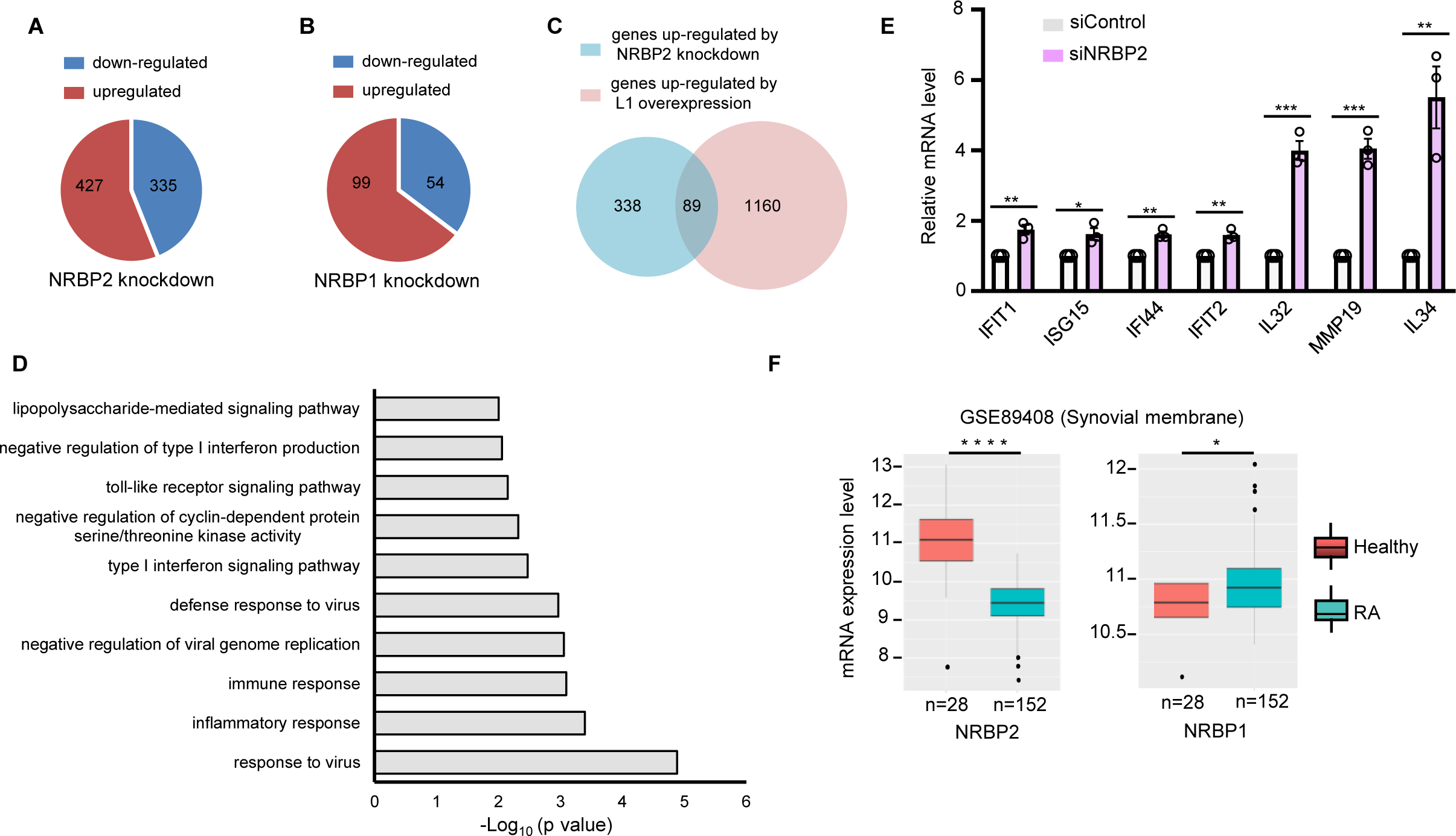
NRBP2 represses innate immune response and NRBP2 mRNA level is negatively correlated with occurrence of RA. **(A and B)** Pie charts illustrating the numbers of genes displaying significantly altered expression levels (fold change ≥ 1.5 and FDR < 0.05) following the knockdown of either NRBP2 **(A)** or NRBP1 **(B)** in HeLa cells. **(C)** Venn Diagram showing overlapping genes upregulated by NRBP2 knockdown in HeLa cells and genes activated upon L1 overexpression in RPE cells. p < 2.081e-25. The p-value was calculated using http://nemates.org/MA/progs/overlap_stats.html. to determine the statistical significance of the overlap between the two groups of genes. **(D)** GO term analysis for the overlapping genes in **(C)** with the online software DAVID. Pathways exhibiting an enrichment with -log_10_p-value >2 are presented. **(E)** qRT-PCR to confirm activation of selected innate immunity genes upon NRBP2 knockdown in HeLa cells. N = 3 biological replicates. **(F)** mRNA levels of NRBP1 and NRBP2 in samples obtained from both healthy individuals and RA patients. Data is taken from Autoimmune Diseases Explorer (https://adex.genyo.es). See also Table S2.

Improper activation of innate immune responses is the cause of some autoimmune diseases and mutations in genes responsible for L1 suppression have often been found in autoimmune diseases.^48,50-52^ NRBP2 mutations have not been reported to be linked to the development of autoimmune diseases. However, from the Autoimmune Diseases Explorer database (https://adex.genyo.es/),^53,54^ we observed a decreased expression of NRBP2 mRNA level within the synovial membrane samples obtained from RA patients (**Figure 6F**). This correlation suggests that NRBP2 might have a protective role against autoimmune diseases such as RA and this could be due to inhibiting retroelements.

### *NRBP1/2* emerged in the early vertebrate lineage by gene duplication and underwent divergent evolution

Invertebrate model organisms, such as *Caenorhabditis elegans* (*C. elegans*) and *Drosophila melanogaste*r have a single *NRBP1/2* homolog, whereas humans and mice have a set of two paralogous genes residing on two different chromosomes. Our data so far suggest human NRBP2 as an inhibitor of NRBP1. To understand how this regulatory relationship emerged, we aimed to trace the evolutionary history of this gene duplication event that led to the observed functional differentiation.

We obtained a set of 2,065 *NRBP* homologs with Blast searches in the UniProt database using human NRBP1 and NRBP2 as queries. We accepted only proteins that, when compared in a second ‘reverse’ Blast search against the human proteome, identified either human NRBP1 or NRBP2 as top-ranking hits. When we applied the criteria for the identification of pseudokinases, we learned that all the 2,065 *NRBP* homologs encode pseudokinases.^55^ Subsequently, we used maximum likelihood to infer the phylogeny of all identified *NRBP* homologs. The resulting phylogenetic analysis recovered 670 *NRBP2* as well as 933 *NRBP1* orthologues as separate monophyletic groups with perfect bootstrap support. The two clades are sister groups, and their branching point was also recovered with perfect bootstrap support, suggesting that the data are highly reliable. The remaining genes are separated from *NRBP1* and *NRBP2* and comprise all invertebrate *NRBP* (**Figure 7A**).

**Figure 7.**
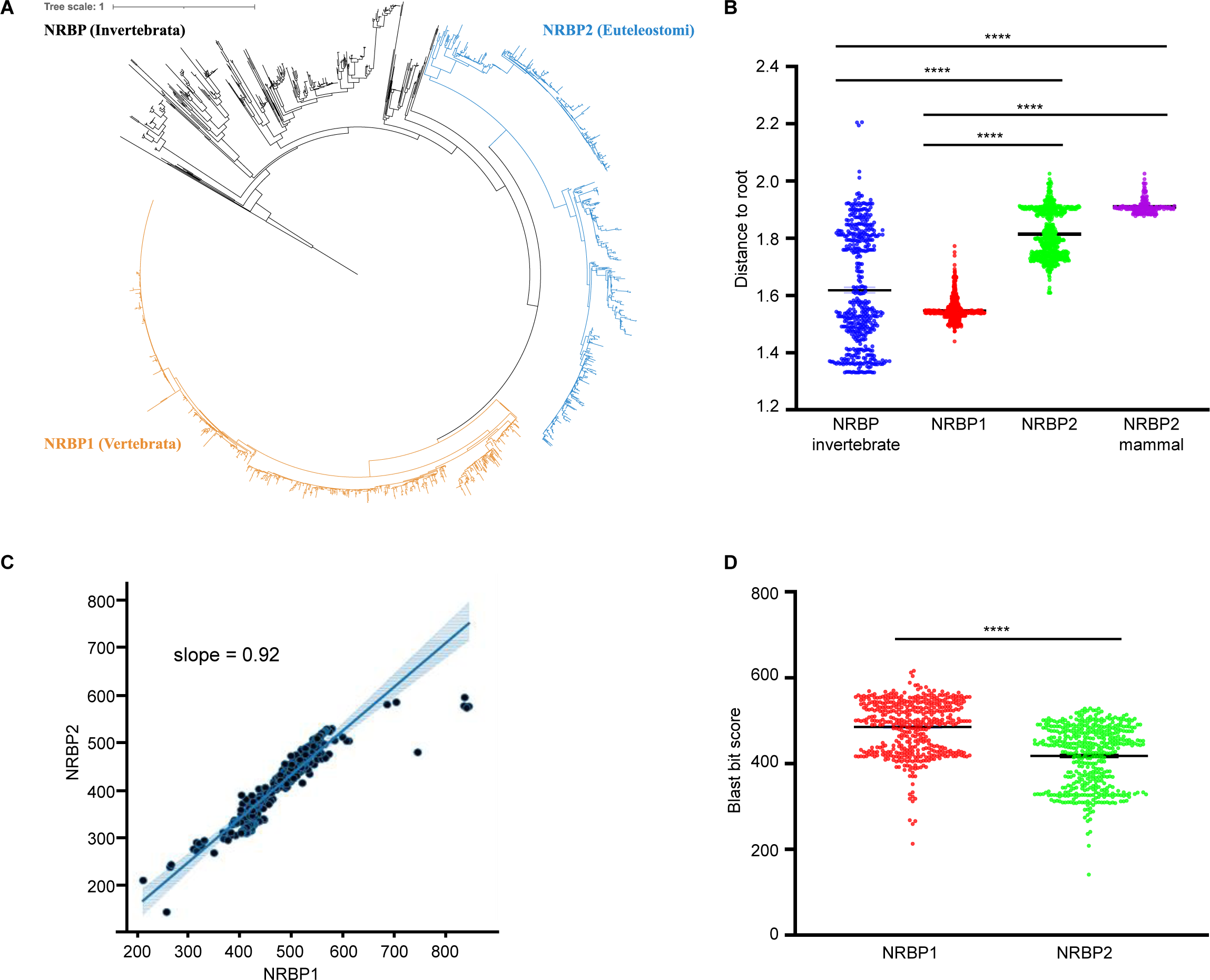
Phylogeny of *NRBP1/2* homologs generated using maximum likelihood. (A) Maximum likelihood phylogeny of the IQ-TREE analysis comprising 2,065 quality-filtered NRBP proteins belonging to 663 species and including 417 amino acid positions. Orthologues of human *NRBP2* are recovered as a monophyletic group with perfect bootstrap support (blue). All *NRBP2* belong to the Euteleostomi. Its sister clade contains all orthologues of human *NRBP1* derived from vertebrate taxa (orange) and has perfect bootstrap support as well. The *NRBP* sequence of the sponge *Amphimedon queenslandica* was chosen as outgroup taxon. Branch lengths are amino acid substitutions per site. (B) The distance from leaf to root was determined in **(A)** for all NRBP proteins. The NRBP2 clade accumulated considerably more amino acid substitutions than the NRBP1 clade and the average invertebrate NRBP protein. A consequence of this is that all invertebrate NRBP proteins are more similar to human NRBP1 than they are to human NRBP2. (C) Scatter plot of BlastP bit scores resulting from 462 comparisons of invertebrate NRBP either to human NRBP1 or to human NRBP2. For all value pairs, the bit score for the NRBP1 comparison is higher than that of the respective comparison to NRBP2. (D) Plot and paired t-test for the values described in **(C)** results in a p-value of 1.5 × 10^-202^. This indicates that human NRBP1 is more similar to invertebrate NRBP than human NRBP2 is. See also Figure S8.

In addition, the respective phylogenetic lineage information of the 2,065 proteins (including human NRBP1 and NRBP2) revealed that *NRBPs* are present in the earliest metazoan clades, including the early-branching Porifera (sponges) and Placozoa. Non-metazoan eukaryotes, such as choanoflagellates, fungi, amoebozoans, plants, bacteria and archaea, do not have *NRBPs* (**Figure S8A**). While *NRBP1* orthologues are placed within the Vertebrata (vertebrate) category, all the *NRBP2* orthologues belong to the Euteleostomi (bony vertebrates), a sub-clade of the vertebrates (**Figure S8B**).

Moreover, by analyzing the leaf to root distances in the phylogenetic tree **(Figure 7A)**, we learned that the proteins comprised in the *NRBP2* clade accumulated considerably more amino acid substitutions than those comprised in the *NRBP1* clade and the average invertebrate *NRBP*. The mammalian *NRBP2* forms a consistent sub-clade, exhibiting an even higher rate of amino acid substitutions (**Figure 7B**). This is consistent with our Blast results, in which all invertebrate *NRBPs* produced higher bit scores when compared to human *NRBP1* than to human *NRBP2* (**Figure 7C, D**).

Taken together, these observations support a scenario in which *NRBP1/2* originated from a single gene duplication event very early in the vertebrate lineage. While the more conserved *NRBP1* probably maintained the original *NRBP* functions, *NRBP2* may have evolved to serve new functional niches, such as regulating NRBP1 activity by targeting it to degradation.

## Discussion

Gene duplication plays a crucial role in driving functional innovation throughout the course of evolution. Proteins encoded by paralogous genes often exhibit redundant functions, and in some cases, they may even display antagonistic activities. This antagonism is primarily attributed to the competition for common binding partners between the gene products of the functional and loss-of-function copies.^4^ A detailed dissection of such a competitive mechanism has explained the antagonistic roles of UPF3A and UPF3B in nonsense-mediated decay machinery.^4^ In our study, we present evidences supporting the idea that the antagonistic relationship between paralogs can be established by directing the degradation of the precursor gene product through the action of the later duplicate product. Our findings suggest that this acquired ability to regulate the precursor has been evolutionarily favoured, probably through improved flexibility in increasingly complex biological systems.

Instead of competing for ORF1 association, we found the inhibitory role of NRBP2 on L1 relies on presence of NRBP1. This is achieved by targeting NRBP1 protein for degradation, probably through ubiquitin–proteasome system (UPS). NRBP1 is known to function in the Elongin B/C E3 ubiquitin ligase complex to facilitate the ubiquitination and decay of specific substrates such as BRI2 and BRI3.^11^ NRBP1 degradation upon NRBP2 overexpression is, however, independent of Elongin B/C. Our confocal imaging reveals that the degradation of NRBP1 is confined primarily to the perinuclear region of cells, likely due to the presence of the relevant UPS components in this subcellular area, such as on or near the ER membrane. This suggest that the inhibitory role of NRBP2 on NRBP1 may be limited to NRBP1 functions specifically associated with perinuclear region. However, for functions of NRBP1 occurring on the plasma membrane, such as regulating signal transduction, NRBP2 may not necessarily deactivate NRBP1. Therefore, identification of the involved E3 ligase would be fundamental to further dissecting functional interaction between NRBP1 and NRBP2. In addition, future work should address which part of the UPS is affected by NRBP2: Does it modulate the ubiquitination/deubiquitination of NRBP1 and its recognition by E3 ligase/deubiquitinase, or other early steps of ubiquitin conjugation? How could binding of NRBP2 to NRBP1 result in its degradation by UPS? NRBP1 is known to form homodimers to function in the Elongin B/C complex. If the ancestral NRBP also forms a homodimer, NRBP2 may have the capacity to form heterodimers with NRBP1 immediately after the gene duplication event, and the capability is possibly conserved during evolution. We propose that the possible heterodimer might render NRBP1 as the substrate of UPS. NRBP proteins are named for the presence of NRB motifs. Surprisingly, our study of the NRBP1/2 interactome does not uncover any nuclear receptors as their interactors. Instead, our biochemical assays highlight the significant role of NRB in facilitating both homo- and heterodimerizations of NRBP proteins. Currently, we cannot definitively determine whether NRB directly participates in dimerization or if its presence is essential for proper protein folding to enable dimerization.

The fact that the C-terminal half of NRBP1 also destabilizes the full-length NRBP1, indicating that presence of the N-terminal halves of both single NRBP1 molecules in the homodimer may prevent its detection by UPS. In contrast, the N-terminal half of NRBP2 cannot fulfil this protective role, suggesting a perturbation of certain functional motifs of amino acids during evolution that converted NRBP2 to an inhibitor of NRBP1. Therefore, although the C-terminal half of NRBP2 is necessary and sufficient for NRBP1 degradation, the critical motifs discriminating these two pseudokinases probably exist in their N-terminal halves. Why does lack of the N-terminal half of NRBP1 in the homodimer trigger its decay? One possible scenario is that the sites for ubiquitin conjugation or E3 ligase binding might then be exposed to E3. Or the UPS just recognizes it as a misfolded and defective protein that fails the protein quality control. Elucidating the ubiquitin site responsible for NRBP1 degradation would provide a valuable hint to answer these questions. In addition, we anticipate that similar regulatory relationship might also exist among other paralog encoded proteins that function as homo-or heterodimers.

The paralogs *NRBP1* and *NRBP2* in this study encode two highly conserved pseudokinases which, according to our sequence analysis, have lost the ability to phosphorylate proteins at the dawn of their emergence. The wide existence of NRBPs in multicellular animals indicate that they might execute important biological functions. A previous genetic screen by us had revealed participation of the *C. elegans* NRBP, HPO-11 in tumorigenesis.^56^ In addition, numerus recent works suggested regulatory roles of human NRBP1 and NRBP2 in tumor biology, as either tumor suppressor or activator. These reports are, however, mostly descriptive without considerable mechanistic details. Our MS interactome analysis provides an unbiased insight into potential molecular functions of NRBP1/2. In addition to the known interactors of the Elongin B/C E3 ligase complex, interactors of NRBP1/2 are mostly enriched in RNA-binding proteins and those in transcriptional regulation, implicating that these two pseudokinases might be involved in RNA- and DNA-related processes (**Table S1**). Here we suggest NRBP1 and NRBP2 as novel regulators of L1 retrotransposon, probably through influencing L1 RNP integrity. Remarkably, the emergence of *NRBP2* in the Euteleostomi coincided with a switch of the L1 ORF1 from a ‘Type I’ to a ‘Type II’. It was reported that vertebrates have ‘Type II’ ORF1, whereas other animal and plant taxa have ‘Type I’ ORF1.^57^ However, when we analysed the phylogenetic distribution of ‘Type II’ ORF1, which is characterized by PFAM domain PF02994,^58^ we recognized that ‘Type II’ ORF1 is actually absent from non-euteleostomian vertebrates. 4404 of 4407 (99.9%) of the metazoan reports of PF02994 in the PFAM database belong to the Euteleostomi, and none are found in Agnatha and Chondrichthyes, which also lack *NRBP2* (**Figure 7B**). The strict correlation of the appearance of *NRBP2* with the switch of L1 transposon to the Type II ORF1 opens the possibility that *NRBP2* might have co-evolved to control this new variant of L1.

Although both NRBP1 knockdown and NRBP2 overexpression result in enrichment of ORF1 in certain cytoplasmic foci, these foci are not identical. The ORF1 foci upon NRBP1 knockdown were smaller and lack of the SG marker protein G3BP1, those induced by NRBP2 overexpression were larger and G3BP1 positive. Since NRBP2 overexpression triggers formation of ORF1 foci and inhibit L1 activity independently of G3BP1, these ORF1 enriched foci are probably not SGs, or are special G3BP1-independent SGs. NRBP2 overexpression could also possibly result in certain cellular stress to induce such SG-similar structures that contain many RNA-binding proteins, including ORF1, in addition to disassociating ORF1 from L1 mRNA. Several of our identified NRBP1 interactors, such as G3BP1 and MOV10, also trigger translocation of both ORF1 protein and L1 mRNA into SGs to inhibit L1, arguing for a quite different strategy used by NRBP1 to regulate L1.^40,59^ One possible scenario could be that NRBP1 acts as an adaptor protein to facilitate assembly of L1 RNP complexes, and lack of NRBP1 could also impair incorporation of the other host factors into L1 RNPs.

The activation of retrotransposons not only induces genome instability, influencing tumor development, but also triggers an innate immune response, contributing to autoimmune diseases. While NRBP2 is recognized as a tumor suppressor, NRBP1 exhibits dual roles, acting as either a tumor suppressor or activator depending on the tumor type. Our correlation analysis additionally indicates that NRBP2 serves as a potential safeguard against autoimmune diseases. Our finding reveals an antagonistic regulation of L1 retrotransposon by NRBP1 and NRBP2, holding potential clinical implications. Specifically, diseases arising from aberrant L1 activation might be treated with NRBP1 inhibitors or NRBP2 activators. In summary, our discovery that NRBP2 targets NRBP1 for degradation offers profound insights into evolutionary biology, retrotransposon regulation, and therapeutic implications in medicine.

## Acknowledgements

We thank the ProteomeXchange Consortium for providing globally coordinated standard data submission. We thank the staff of Life Imaging Center (LIC) at the University of Freiburg for providing access to their microscopy resources and for their outstanding technical support. We thank China Scholarship Council (CSC) for providing financial support to S.C. Work in the lab of B.W. was supported by the Deutsche Forschungsgemeinschaft (DFG, German Research Foundation), transregional collaborative research grant TRR130. This work was funded by grants from DFG (SFB1381) and Germany’s Excellence Strategy (CIBSS-EXC-2189-Project ID 8390939984) to R.B.

## Author contributions

W.Y. and W.Q. conceived/designed the experiments, analysed data, wrote the manuscript. W.Y. and S.C performed most of the experiments. S.U. performed the experiments in Figure 4E, 6E, S5E and S5I. D.O. did IP in Figure S1B. P.E. and A. Trentino contributed to cloning. J.S., K.F., A.Thien and B.W. contributed to mass spectrometry experiments and data analysis. T.H. provided material support. T.S. and E.S. performed phylogenetic analyses and wrote the respective sections of the manuscript. R.B. provided funding for the project and conceptional suggestions for its execution and participated in writing the manuscript.

## Materials and Methods

### Cell culture

HeLa alpha Kyoto^60^ and HEK293T were cultivated in Dulbecco’s modified Eagle’s medium (DMEM) with 4.5 g/L glucose (PAN), 10% fetal bovine serum (FBS, Gibco) and 3 mM L-glutamine (Gibco). MCF-7 cells (ACC115, DSMZ) were cultivated in RPMI 1640 (PAN), supplemented with 10% FBS (Gibco) and 3 mM L-glutamine (Gibco).

### Plasmids

The L1 reporter, pCMV-L1-neo^TNF^ (L1-neo) was generated in ^39^. pCMV-L1-Flag-neo^TNF^ (L1-Flag-neo, pBY4215) was generated by inserting an in-frame Flag tag to the C-terminus of ORF1 in L1-neo plasmid. To construct HA-NRBP1 (pBY4258), HA-NRBP1-N (pBY4278), HA-NRBP1-C (pBY4282), HA-NRBP1-C-dNRB (pBY4287), HA-NRBP2 (pBY4216), HA-NRBP2-C (pBY4271) and HA-NRBP2-C-dNRB (pBY4288), the corresponding coding sequences (CDS) were amplified by PCR and inserted into pRK5-HA vector with the *EcoRI* / *HindIII* sites. To generate NRBP1-myc (pBY4237), NRBP1-N-myc (pBY4281), NRBP1-C-myc (pBY4283), ΔLCR-NRBP1-myc (pBY4254), NRBP2-myc (pBY4243), NRBP2-N-myc (pBY4245) and NRBP2-C-myc (pBY4218), NRBP2-dNES-myc (pBY4233), NRBP2-dNLS-myc (pBY4238), NRBP2-dBC-myc (pBY4244), NRBP2-dNRB-myc (pBY4250) and NRBP2-dDimer-myc (pBY4255), LCR-NRBP2-myc (pBY4259), the individual CDS was amplified with primers containing an in-frame stop codon to terminate the N-terminal HA tag in the pRK5-HA vector and the Myc tag was fused at the C-terminus of the CDS. The PCR products were inserted into the *EcoRI* / *HindIII* sites of pRK5-HA. To generate NRBP1-Flag (pBY4275), NRBP1 CDS was amplified with primers containing an in-frame stop codon to terminate the N-terminal HA tag in the pRK5-HA vector and Flag tag was added at the C-terminus of the CDS. The PCR product was inserted into the *EcoRI* / *HindIII* sites of pRK5-HA. Myc-DDK-NRBP1 (pBY4284) and Myc-DDK-NRBP2 (pBY4285) were generated by inserting NRBP1 and NRBP2 PCR products between the *AscI* / *NotI* sites of the pCMV6-AN-Myc-DDK vector (Origene). ORF1-Flag (pBY4211) was constructed by inserting ORF1-Flag PCR product into the *BamHI* / *EcoRI* sites of pCDNA3.1+ plasmid. shControl (RHS4743), shNRBP1-1, shNRBP1-2, shNRBP2-1 and shNRBP2-2 are from Dharmacon. Target sequences are shown in **Table S3**.

### Immunoprecipitation and in-gel digestion for Mass Spectrometry (MS)

MCF-7 shControl cells, transduced with shControl-expressing viruses, were used for the NRBP1/2 interactome analysis, as NRBP2 specific interactors were also investigated via comparing co-immunoprecipitated proteins in the shControl with NRBP1 knockdown cells. The NRBP2-specific interactome is beyond the scope of this work and therefore not shown. To perform the immunoprecipitation, cells were lysed in CHAPS-based buffer containing 40 mM HEPES (PH7.5), 120 mM NaCl, 0.3 % CHAPS supplemented with Complete Protease Inhibitor Cocktail (11697498001, Roche), Phosphatase Inhibitor Cocktail 2 (P5726, Sigma) and Cocktail 3 (P0044, Sigma). Proteins were immunoprecipitated with NRBP1/2 antibody (21549-1-AP, Proteintech) or normal rabbit IgG (sc-2027, Santa Cruz) coupled with Dynabeads™ Protein G (10009D, Thermo Fisher). The precipitated proteins were separated by NuPAGE® Novex® 4–12% Bis-Tris gels (NP0322BOX, Invitrogen). In-gel digestion with trypsin (Promega, Mannheim, Germany) was performed essentially as described in ^61^. Proteins were destained using 50% ethanol in 10 mM ammonium bicarbonate. For reduction of disulfide bonds and subsequent alkylation, 5 mM tris (2-carboxyethyl) phosphine (10 min at 60 °C) and 100 mM 2-chloroacetamide (15 min at 37 °C) were used, respectively.

### High-Performance Liquid Chromatography and MS

LC-MS analysis was performed either on an UltimateTM 3000 RSLCnano system coupled to an LTQ Orbitrap XL (two replicates) or a Velos Orbitrap Elite instrument (one replicate). All instruments are from Thermo Fisher Scientific, Bremen, Germany. On both HPLCs a binary solvent system was used with solvent A consisting of 0.1% formic acid and 4% DMSO and solvent B consisting of 48% methanol, 30% acetonitrile, 0.1% formic acid and 4% DMSO. The HPLC coupled to the LTQ Orbitrap XL was equipped with two PepMapTM C18 μ-precolumns (ID: 0.3 mm × 5 mm, 5 µm, 300 Å Thermo Fisher Scientific) and an AcclaimTM PepMapTM analytical column (ID: 75 μm × 250 mm, 3 μm, 100 Å, Thermo Fisher Scientific). Samples were washed and concentrated for 5 min with 0.1% trifluoroacetic acid on the pre-column. A flow rate of 0.250 µl/min was applied to the analytical column and the following gradient was used: 1% B to 30% B in 34 min, to 45% B in 12 min, to 70% B in 14 min, to 99% B in 5 min, 99% B for 5 min and decreased to 1% B in 1 min. The column was re-equilibrated for 19 min with 1% B.

The HPLC coupled to the Velos Orbitrap Elite instrument was equipped with two nanoEase™ M/Z Symmetry C18 trap columns (100Å pore size, 5 µm particle size, 20 mm length, 180 µm inner diameter) and a nanoEase™ M/Z HSS C18 T3 analytical column (250 mm length, 75 µM inner diameter, 1.8 µM particle size, 100 Å pore size), all from Waters Corporation, Milford, MA. The trap columns were operated at a flowrate of 10 µl/min and the analytical column at 300 nl/min. Peptide samples were pre-concentrated on the trap column using 0.1% trifluoroacetic acid for 5 min before switching the column in line with the analytical column. Peptides were separated using a multi-step gradient. Over the course of 65 min, the percentage of solvent B increased from 3% to 55%, followed by an increase to 80% B in 5 min. The column was eluted for another 5 min with 80% B before returning to 3% B in 4 min. The column was re-equilibrated for 21 min with 3% B.

The MS instruments were operated with the following parameters: 1.5 kV spray voltage, 200 °C capillary temperature. Orbitrap mass range on both instruments m/z 370 to 1,700. For the LTQ Orbitrap XL the resolution at m/z 400 was 60,000, automatic gain control 5x10^5^ ions, max. fill time, 500 ms. For the Velos Orbitrap Elite the resolution at m/z 400 was 120,000, automatic gain control 1x10^6^ and the maximum ion time 200 ms. A TOP5 (LTQ Orbitrap XL with automatic gain control 10,000 ions, max. fill time 100 ms) or a TOP 25 (Velos Orbitrap Elite with automatic gain control 5,000, max. fill time 150 ms) method was applied for collision-induced dissociation of multiply charged peptide ions. On both instruments the normalized collision energy was 35% and the activation Q 0.250. Dynamic exclusion was set to 45s.

### MS Data analysis

Raw files were searched with MaxQuant version 2.4.9.0^62,63^ against the *homo sapiens* Uniprot reference proteome (ID: UP000005640; 20594 protein entries; October 2022).

Essentially, default settings were used in MaxQuant. Trypsin/P was used as proteolytic enzyme and up to two missed cleavages were allowed. A 1% false discovery rate was applied to both peptide and protein lists. Methionine oxidation and N-terminal acetylation were set as variable and carbamidomethylation as fixed modifications. The minimum number of unique peptides was set to 1. Label-free quantification^64^ was enabled, with a minimum ratio count of two and the option ‘require MS/MS for label-free quantification (LFQ) comparisons’ enabled.

For data analysis, the proteingroups.txt file of Maxquant was used and loaded into Perseus 2.0.10.0.^65^ Entries for reverse and contaminant hits as well as proteins only identified by site were removed from the analysis. LFQ intensities were log_10_-transformed. Only protein groups with at least six reported LFQ intensities were considered for further analysis. Missing values were imputed from normal distribution using the following settings in Perseus: width 0.5 and down shift 1.7. A one-sided two sample t-test was performed with a valid value (non-imputed) filter set for 2 out of 3 in the NRBP1/2 IP replicates. Proteins with a p-value below 0.05 and a student’s t-test difference of at least 5 were considered as significantly enriched candidates.

The mass spectrometry proteomics data have been deposited to the ProteomeXchange Consortium (http://proteomecentral.proteomexchange.org) via the PRIDE partner repository^66^ with the dataset identifier PXD051452.

### Co-immunoprecipitation (Co-IP)

For the Co-IP experiments, cells were transfected with the respective plasmids and collected 48 hours after transfection. The cells were washed twice with PBS and lysed in the CHAPS-based buffer as stated above, supplemented with Complete Protease Inhibitor Cocktail, Phosphatase Inhibitor Cocktail 2 and Cocktail 3. The cell lysates were pre-cleared with Dynabeads™ Protein G for 30 minutes and then incubated with specific antibodies coupled with the protein G magnetic beads for 2-3 hours. After washing the beads with CHAPS-based buffer, the proteins were eluted by boiling at 95 °C for 5 min in Laemmli Sample Buffer (containing 10% glycerol, 1% beta-mercaptoethanol, 1.7% SDS, 62.5 mM Tris pH 6.8, and bromophenol blue).

### Cell lysis and Western blot

Cells were washed twice with PBS and then lysed with RIPA buffer (150 mM NaCl, 50 mM Tris pH 8.0, 1 % NP40, 0.5 % sodium deoxycholate, and 0.1 % SDS), supplemented with Complete Protease Inhibitor Cocktail, Phosphatase Inhibitor Cocktail 2 and Cocktail 3 for 10 minutes before centrifugation. The protein concentration was measured using Bio-Rad Protein Assay Dye Reagent Concentrate (#5000006) and adjusted to the same level. The cell lysates were mixed with Laemmli Sample Buffer and heated for 5 minutes at 95 °C. Cell lysates were loaded onto sodium dodecyl-sulfate polyacrylamide gel electrophoresis (SDS-PAGE) gels and the proteins were transferred to polyvinylidene difluoride (PVDF) membranes. The membranes were blocked with 5% bovine serum albumin (BSA) diluted in Tris-Buffered Saline with Tween 20 (TBST) buffer for 1 hour at room temperature and then incubated with primary antibodies overnight at 4 °C. Membranes were washed with TBST buffer and incubated with the corresponding horseradish peroxidase (HRP)-conjugated secondary antibodies for 1-2 hours. Pierce ECL Western Blotting Substrate (32209) or SuperSignal West Femto Maximum Sensitivity Substrate (34095) were used to detect the protein signals. The chemiluminescence signal was captured using a LAS-4000 camera system.

### siRNA knockdown

ON-TARGET plus SMARTpool siRNAs directed against NRBP1(L-005356-00), NRBP2 (L-005340-02), ELOB (L-012376-00), ELOC (L-010541-00) and Non-targeting control (siControl) siRNA (D-001810-10) were purchased from Dharmacon. HeLa cells were transfected with a final concentration of either 20 nM siRNA (for NRBP1 and NRBP2 knockdown) or 10 nM siRNA (for ELOB and ELOC knockdown) in two consecutive days using the Lipofectamine RNAiMAX Transfection Reagent (Thermo Fisher Scientific) according to the manufacturer’s instructions. For NRBP1 and NRBP2 knockdown, cells were collected for analysis two days after the second siRNA transfection. For **Figure S7B, C**, cells were transfected with plasmids one day after the last siRNA transfection. Cells were collected 28 hours after the plasmid transfection.

### shRNA knockdown

The pTRIPZ doxycyclin-inducible shRNA constructs targeting NRBP1 and NRBP2 or the non-targeting control sequence (shControl) were obtained from Dharmacon. Information of the target sequence is given in **Table S3**. Viral particles were produced using the Trans-Lentiviral shRNA Packaging mix (Horizon Discovery) according to the manufacturer’s protocol. HeLa and MCF-7 cells were transduced with the viral particles in the presence of 8 µg/mL polybrene. Puromycin (2 µg/mL) selection was carried out 48 hours post-transduction for 7 days. Expression of the shRNA was induced with 2 µg/mL doxycycline for indicated days as mentioned, respectively.

### Generation of G3BP1 knockout cell lines

To generate G3BP1 CRISPR/Cas9 knockout HeLa cell lines, three target sequences were selected from the human CRISPR knockout pooled library (Brunello, Addgene #73178) and cloned into pSpCas9(BB)-2A-Puro (PX459) V2.0 vector (Addgene # 62988). HeLa cells were transfected with a pool of the three plasmids using Lipofectamine 2000. The pSpCas9(BB)-2A-Puro (PX459) V2.0 vector was also transfected to generate sgControl cells. Twenty-four hours after transfection, cells were selected with puromycin (2 µg/mL) for two days. Monoclonal cell populations were obtained by limiting dilutions. Knockout efficiency was confirmed by Western blot. Target sequences are shown in **Table S3**.

### L1 retrotransposition assay

To test the effect of NRBP1 and NRBP2 overexpression on L1 retrotransposition, HeLa cells were transfected with either L1-neo or pCDNA3.1 together with NRBP1 or NRBP2 expression plasmids. For experiments performed in knockdown cell lines, the inducible cell lines were first treated with 2 µg/mL doxycycline for three days before transfection of the L1-neo or pCDNA3.1 plasmid. Doxycycline was removed until the cells were trypsinized for selection with G418. For experiments carried out in knockout cells, the cells were transfected with only L1-neo or pCDNA3.1. For all experiments, cells were trypsinized 48 h post-transfection, and the same number of cells were re-plated in 6-well plates and selected with G418 (800 µg/ml) (11811031, Thermo Fisher scientific) for 8-14 days. The G418 containing medium was changed every two or three days. To stain the colonies, cells were washed twice with PBS, fixed with methanol for 10 min, and stained with 0.5% crystal violet for 10 min.^67^ The number of G418 resistant colonies was counted using ImageJ software. The relative retrotransposition activity is calculated by dividing the colony numbers in L1-neo by the colony numbers in pCDNA3.1.

### RNA immunoprecipitation (RIP)

MCF-7 shControl and MCF-7 shNRBP1-1 cells were treated with 2 µg/mL doxycycline for 3 days. HeLa cells were transfected with L1-Flag-neo together with HA-NRBP2 plasmids for 2 days. Cells were washed twice with cold DPBS and then lysed in NP-40 Buffer containing 150 mM NaCl, 50 mM Tris pH8.0, 1% NP40, protease inhibitor and RNase inhibitor for 30 minutes. The supernatant was collected by centrifugation at 12,000 rpm for 20 minutes at 4 °C and pre-cleared by incubating with protein G for 30 minutes. The pre-cleared supernatant was then incubated with protein G and ORF1 or Flag antibody overnight at 4 °C. The beads were washed three times and digested by DNaseI for 30 minutes at 37 °C. The reaction was stopped by EDTA, and RNA was extracted using Trizol (15596018, Invitrogen).

### Quantitative Real-Time PCR (qRT-PCR)

Total RNA extraction and DNA digestion were performed using FastGene RNA Premium Kit (FG-81050, Nippon Genetics) unless otherwise specified. cDNA was synthesized with Oligo-dT primer by using Transcriptor High Fidelity cDNA Synthesis Kit (5081963001, Roche). qRT-PCR was performed using Luna Universal qPCR Master Mix (M3003, NEB) running on the Light Cycler 96 System (Roche). Control reactions without reverse transcriptase (no RT) were performed to confirm the absence of contaminating DNA. The data were analyzed by the relative quantification 2-ΔΔCt method.^68^ GAPDH was used for normalization. All qRT-PCR primers used in this study are provided in **Table S3**.

### RNAseq

The RNAseq data were analyzed on the European Galaxy server (usegalaxy.eu). The quality of FASTQ files was checked by FastQC. Cutadapt was used to remove the adapters.^69^ Trimmed reads were aligned to the human genome USCS build hg38 using RNA STAR (v2.7.6a).^70^ The number of reads was counted by FeatureCounts using default parameters.^71^ Differentially expressed genes were identified by EdgeR.^72^ Genes were considered to be significantly differentially expressed when the FDR < 0.05, with log2 fold change ≥ 0.58 (1.5-fold change) for upregulated genes and ≤ -0.58 (1.5-fold change) for downregulated genes. The RNAseq data are available in **Table S2**. RNA-seq data have been deposited in BioProject under ID PRJNA1101872.

### Immunofluorescence and confocal microscopy

Cells seeded on coverslips were washed with PBS, fixed in 4% PFA for 20 minutes, and permeabilized with 0.1% Triton-X100 / PBS for 5 minutes. The cells were then incubated in a blocking solution (3% FBS in PBS) for 20 minutes and with the indicated primary antibodies for 1 hour at room temperature. Subsequently, the cells were incubated with the corresponding Alexa Fluor-coupled secondary antibodies for 40 minutes at room temperature, and the nuclear DNA was stained with Hoechst 33342 (H3570, Sigma). ProLong Gold antifade (P36930, Invitrogen) was mounted on top of the cells. Confocal microscopy was performed using either an LSM-U-NLO or LSM-I-NLO confocal microscope (Carl Zeiss).

### Single-molecule fluorescent in situ hybridization (smFISH)

MCF-7 shControl and shNRBP1-1 cells were treated with 2 µg/mL doxycycline for 3 days before smFISH staining was performed. In total, 48 Stellaris RNA FISH probes, each 20 nt in length targeting L1 mRNA, were produced and purified by Biocat.^73^ The probes were conjugated with Quasar 670 (sequences of the probe sets are available in **Table S3**). ORF1 was stained with the ORF1 antibody (ab230966, Abcam). The smFISH staining was conducted following ‘The Protocol for simultaneous immunofluorescence (IF) + Stellaris RNA FISH in adherent cells’, accessible online (https://www.biosearchtech.com/support/resources/stellaris-rna-fish). Images were captured using a confocal LSM-I-NLO microscope (Carl Zeiss) with Airyscan super-resolution mode.

### Phylogenetic analysis of *NRBP*

To identify homologs of human *NRBP1* and *NRBP2*, we performed BlastP (vs. 2.15.0+) searches for both proteins in UniProtKB/Swiss-Prot and UniProtKB/TrEMBL (release 2024_01) using an E-value cutoff of 1e-28.^74-76^ This threshold was required to prevent the detection of unrelated protein kinase families. Subsequently, we tested the resulting 8,345 sequences with a ‘reverse’ BlastP search in the human proteome (ID: UP000005640). If either NRBP1 or NRBP2 was the top-ranking hit, we kept the respective sequence, otherwise it was rejected. After quality filtering for 80% or higher NRBP1/2 coverage in the BlastP alignments and a further filtering step for the representation of the complete predicted folded region of NRBP1/2 from AlphaFold^44,45^ with a tolerance of 40 amino acid residues, we obtained a set of 2,065 proteins. We obtained the taxonomic information of the respective species using a service from The European Nucleotide Archive (ENA).^77^ In order to conduct phylogenetic analysis, both data sets were merged and sequences were aligned using MAFFT v.7.487 with 1000 iterative refinements.^78^ Alignment masking was performed with trimAl v.1.4.1 by removing all alignment positions containing gaps in 10% or more of the sequences.^79^ Phylogenetic inference was computed with the maximum likelihood method using IQ-TREE v.2.1.2 in conjunction with model selection via ModelFinder to identify the most fitting model of sequence evolution as well as utilizing 1,000 ultrafast bootstraps to assess branch support.^80-82^ Tree evaluation and graphics were done with iTOL v6.9.^83^ Leaf to distances were measured with the Bio.Phylo package using clade labels exported from iTOL.^84^

### Sequence analysis of the pseudokinase domains of NRBP

We predicted the absence of protein kinase activity by scoring three amino acid residues of the ‘catalytic triad’, comprised of the ATP-binding β3-lysine, the catalytic aspartate within the catalytic loop HRDXXXN motif, and the metal binding aspartate of the activation loop DFG motif.^55^ If two of the three amino acid residues did not match the protein kinase consensus, we rated the proteins as pseudokinases. For the identification of the respective amino acid residues, we generated HMMSEARCH (vs. 3.3, http://hmmer.org/) alignments with the PFAM protein kinase domain PF00069 for each sequence.^58^

### Statistics

The presented data includes error bars depicting the standard error of the mean (SEM). Statistical significance was assessed using two-tailed and unpaired Student’s t-tests. The number of replicates is shown in the figures. A P-value <0.05 was considered significant. n.s., not significant. * P < 0.05, * * P < 0.01, * * * P < 0.001, * * * * P < 0.0001.

## Supplemental figure legends

**Figure S1.**
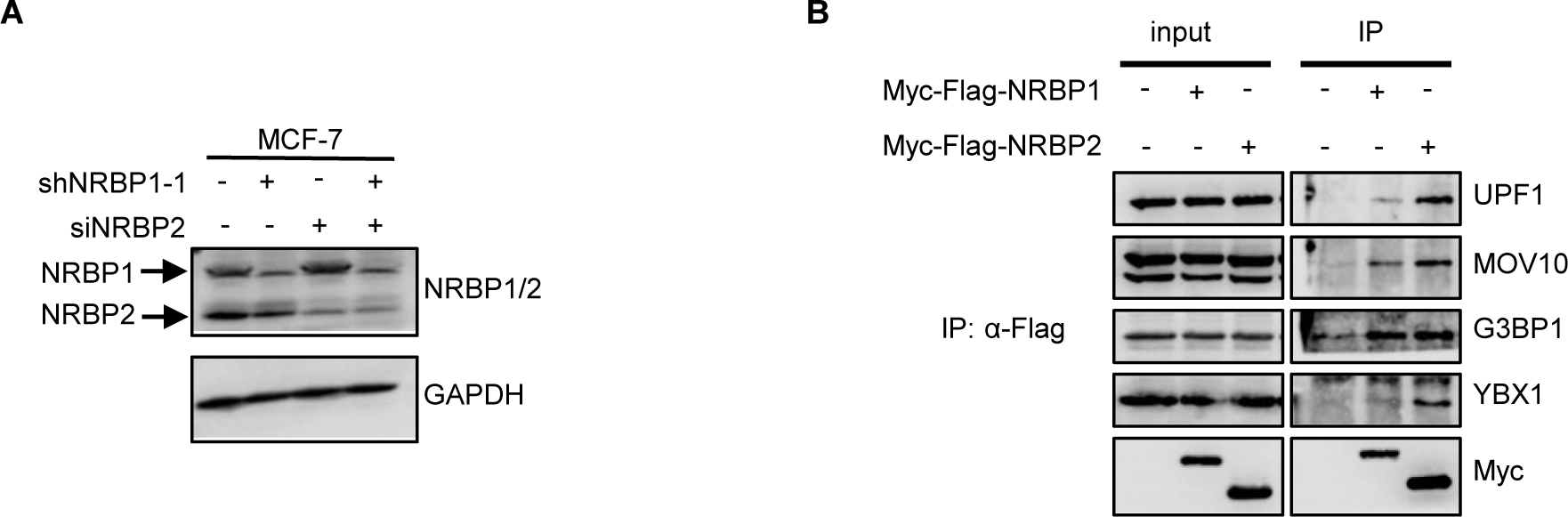
NRBP1 and NRBP2 interact with L1-encoded ORF1 and other ORF1 interactors, related to Figure 1. **(A)** The NRBP1/2 antibody used for MS interactome analysis recognizes both NRBP1 and NRBP2. **(B)** Both NRBP1 and NRBP2 are associated with multiple known L1 interactors or regulators. HEK293T cells were transfected with the indicated plasmids. Flag antibody was used to pull down Myc-Flag-NRBP1 or Myc-Flag-NRBP2, co-precipitated endogenous proteins were detected by using respective antibodies. Myc-Flag-NRBP1 and Myc-Flag-NRBP2 were detected with Myc antibody.

**Figure S2.**
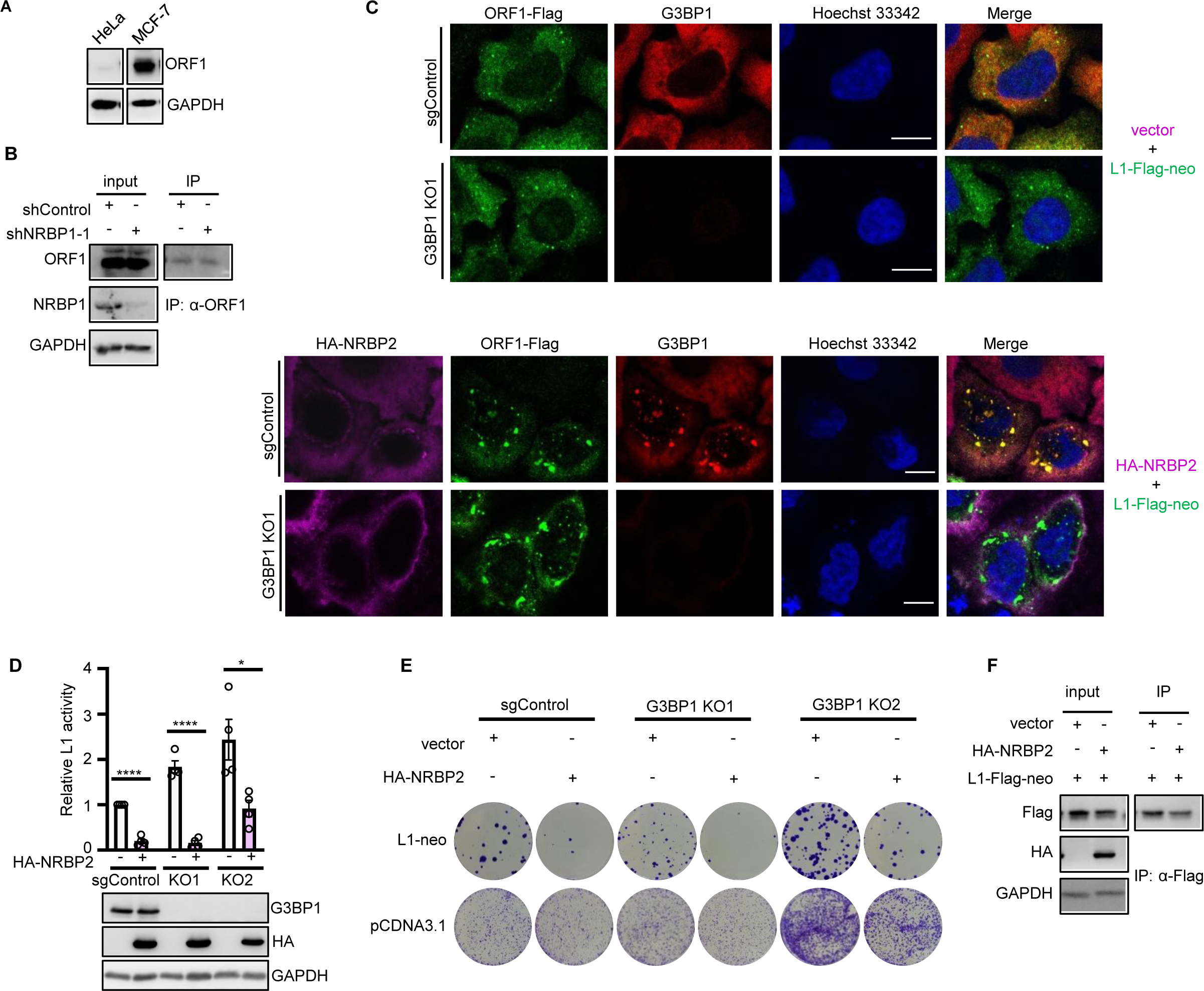
NRBP2-mediated L1 inhibition is independent of G3BP1, related to Figure 2. **(A)** MCF-7 cells show higher ORF1 protein level than HeLa cells. **(B)** Western blot assesses efficiency of NRBP1 knockdown and ORF1 IP for the experiments in Figure 2C. **(C)** Induction of ORF1-Flag enriched foci by NRBP2 overexpression is independent of G3BP1. HA-NRBP2, ORF1-Flag and G3BP1 were stained by HA, Flag and G3BP1 antibodies, respectively. Scale bar 10 µm. **(D)** Inhibition of L1 retrotransposition by NRBP2 is independent of G3BP1. Quantification of the colony assay is shown in the top panel and one representative Western blot result is shown in the bottom panel. N = 4. **(E)** One representative picture of the colony assay depicted in **(D)**. **(F)** Western blot assesses efficiency of HA-NRBP2 transfection and ORF1 IP for the experiments in Figure 2E.

**Figure S3.**
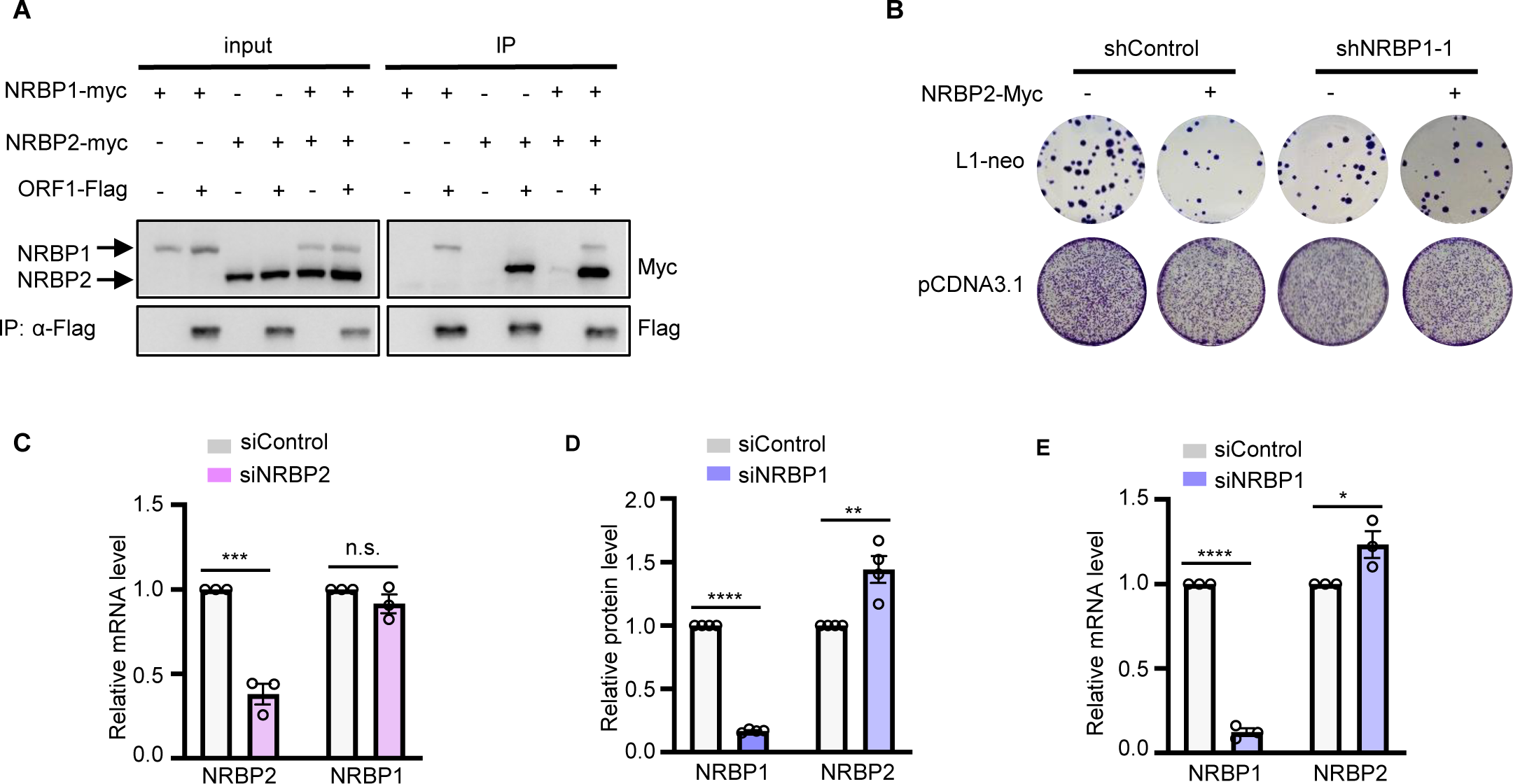
NRBP2 negatively regulates NRBP1 to inhibit L1 retrotransposition, related to Figure 3. **(A)** Overexpressing NRBP1 or NRBP2 does not interfere with binding of their ortholog counterpart to ORF1. Co-IP was performed in HeLa cells by using Flag antibody. **(B)** NRBP1 knockdown abolishes inhibitory effect of NRBP2 on L1 retrotransposition. Shown is a representative picture of the colony assay depicted in Figure 3A. **(C)** NRBP2 knockdown does not significantly increase mRNA level of NRBP1. Shown is quantification of relative mRNA levels of NRBP1 and NRBP2 normalized by GAPDH in HeLa cells. N = 3 biological replicates. **(D)** NRBP1 knockdown increases protein level of NRBP2. Shown is quantification of four replicates in Figure 3C. **(E)** NRBP1 knockdown increases mRNA level of NRBP2. Shown is quantification of relative mRNA levels of NRBP1 and NRBP2 normalized by GAPDH in HeLa cells. N = 3 biological replicates.

**Figure S4.**
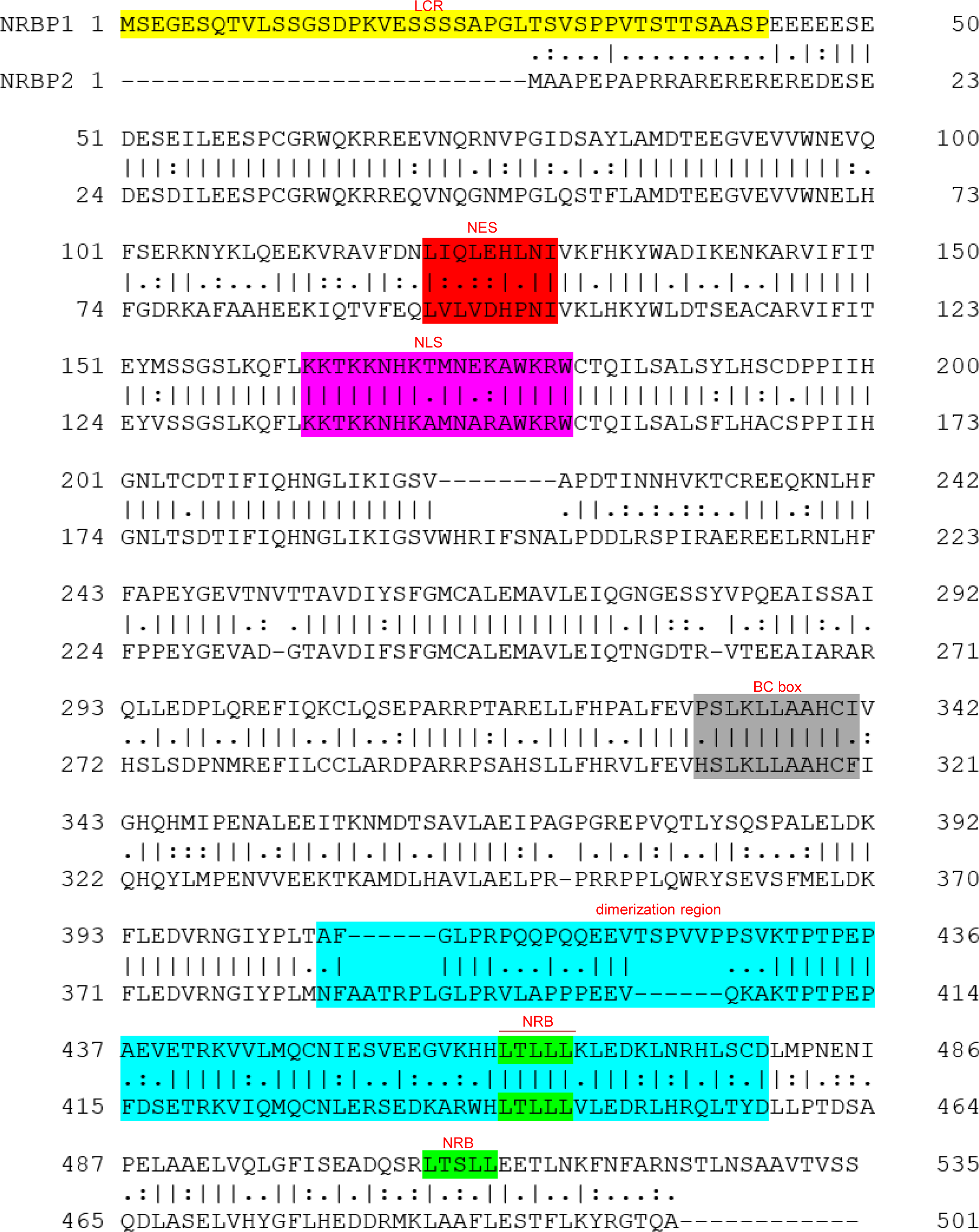
Alignment of the amino acid sequences of NRBP1 and NRBP2, related to Figure 4. Yellow: LCR;^43-45,85^ red: NES; violet: NLS; gray: BC box; blue: dimerization region; green: NRB.^7,9^

**Figure S5.**
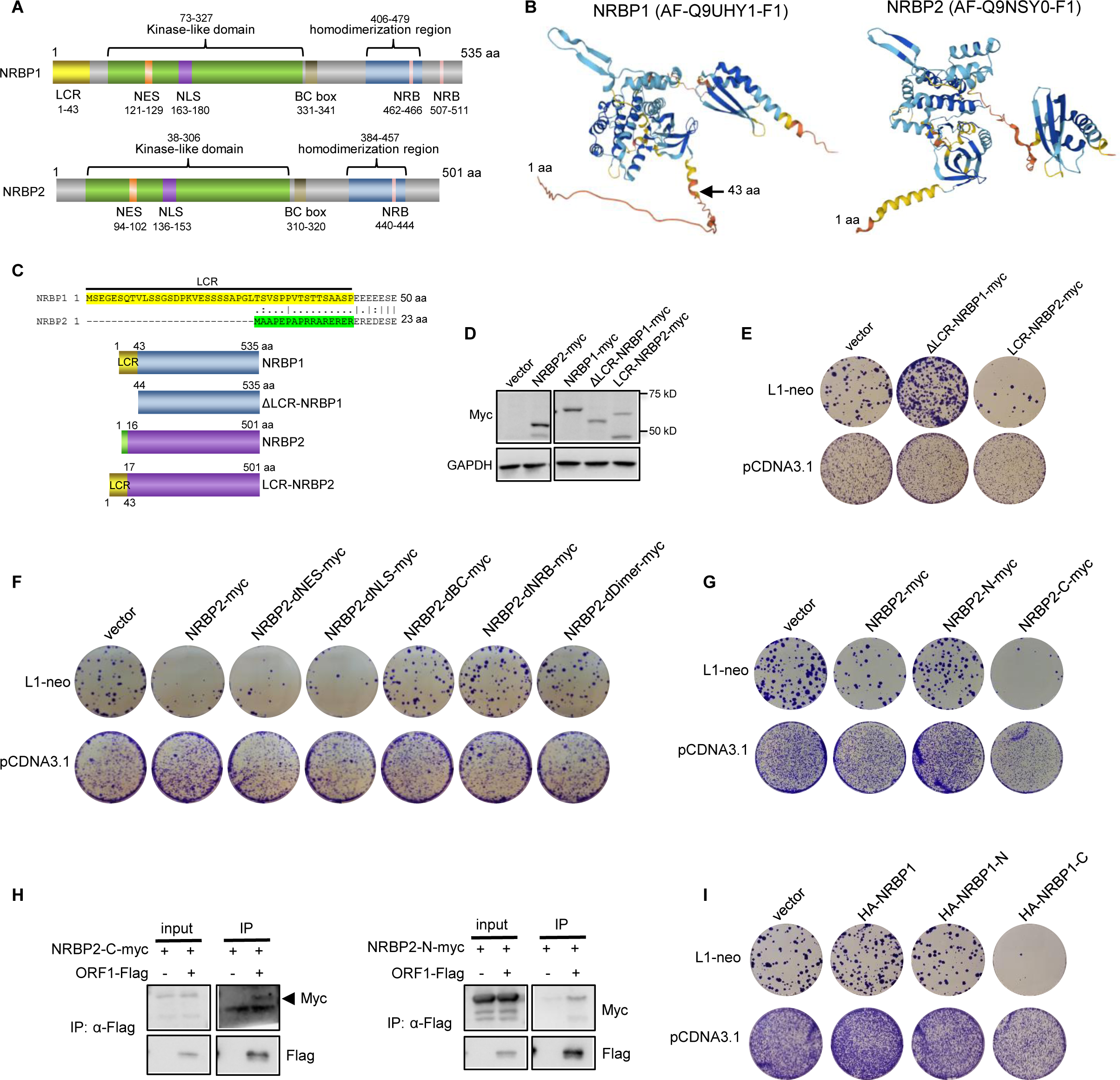
The C-terminal halves of both NRBP2 and NRBP1 negatively regulate L1 retrotransposition, related to Figure 4. **(A)** Diagram illustrations of domains / motifs in human NRBP1 and NRBP2 proteins. **(B)** NRBP1 structure prediction by AlphaFold (AF-Q9UHY1-F1) reveals a potential unstructured N-terminal LCR. **(C)** A simple schematic diagram of NRBP1, NRBP2 and their mutants for the LCR-swapping experiment. **(D)** A representative Western blot to show expression of NRBP1, NRBP2 and their mutants in **(C)** and **(E)**. **(E)** LCR-swapping does not interfere with the regulatory roles of NRBP1 and NRBP2 in L1 mobility. Shown is one representative picture of the colony assay to examine L1 mobility. **(F)** NRB and dimerization region of NRBP2 are essential for its inhibitory role on L1. Shown is one representative picture of the colony assay depicted in Figure 4B. **(G)** The C-terminal half of NRBP2 is necessary and sufficient to inhibit L1. Shown is one representative picture of the colony assay depicted in Figure 4C. **(H)** Both C- and N-terminal halves of NRBP2 interact with ORF1 in HEK293T cells. **(I)** The C-terminal half of NRBP1 functions oppositely to the full-length NRBP1 and inhibits L1. Shown is one representative picture of the colony assay depicted in Figure 4E.

**Figure S6.**
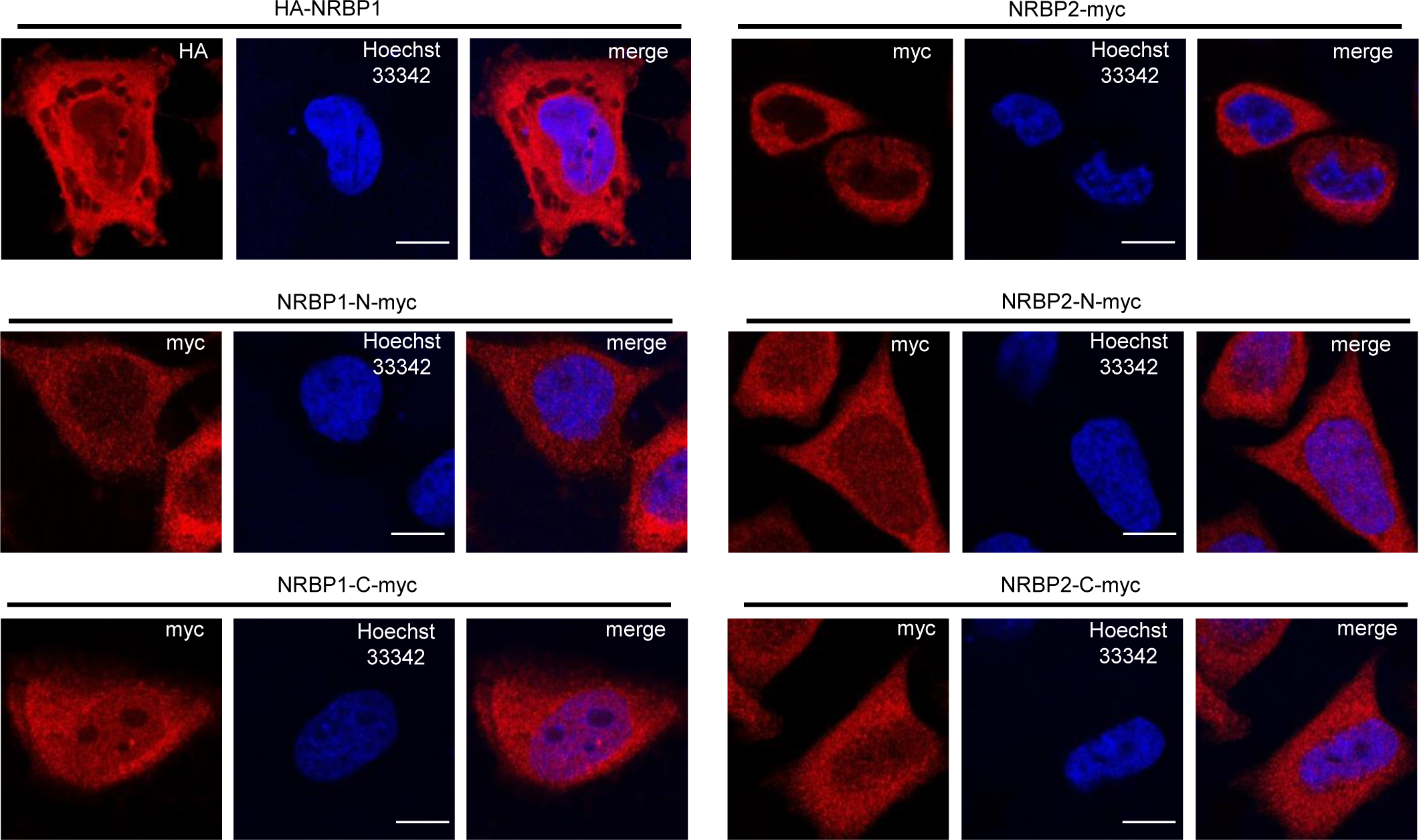
Subcellular localization of overexpressed HA-NRBP1, NRBP2-Myc and their respective N-terminal and C-terminal halves in HeLa cells, related to Figure 4 and 5. Immunofluorescence staining was performed with either HA or Myc antibody. Scale bar 10 µm.

**Figure S7.**
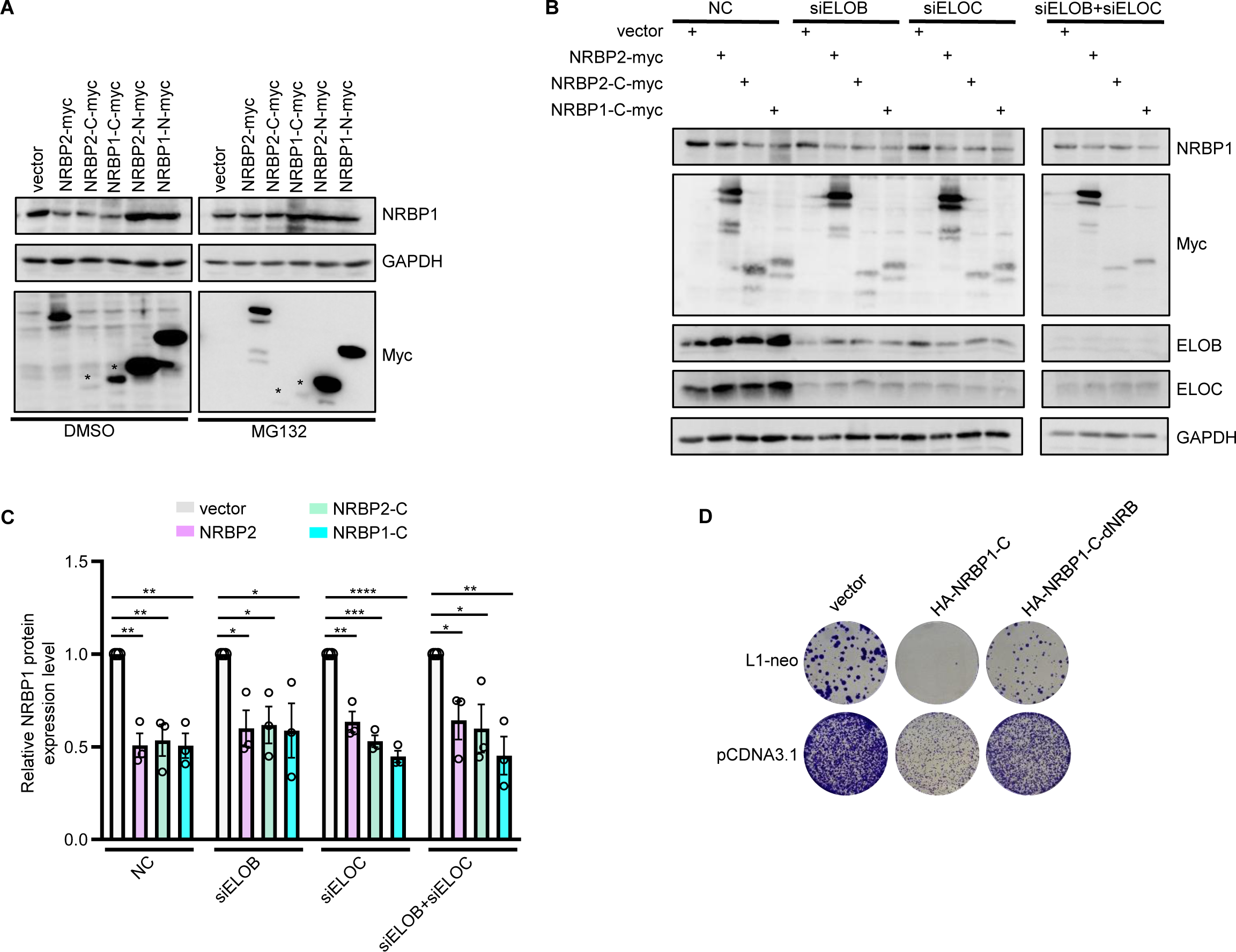
NRBP2 promotes proteasome-mediated NRBP1 decay independent of Elongin B/C E3 ubiquitin ligase complex, related to Figure 5. **(A)** Both the full-length NRBP2 and the C-terminal half of NRBP1 or NRBP2 reduce protein level of full-length NRBP1 in HeLa cells and this could be blocked with MG132 treatment. Shown is one representative Western blot result used for the quantification in the main Figure 5A. **(B)** Both the full-length NRBP2 and the C-terminal half of NRBP1 or NRBP2 promote NRBP1 decay independently of ELOB and ELOC. In line with prior findings, reducing either ELOB or ELOC resulted in a decrease in the protein levels of the other.^86,87^ Shown is one representative Western blot result used for the quantification in **(C)**. **(C)** Quantification of three independent Western blot experiments from **(B)**. **(D)** The NRB motifs in the C-terminal half of NRBP1 are essential to inhibit L1. Shown is one representative picture of the colony assay depicted in the main Figure 5D.

**Figure S8.**
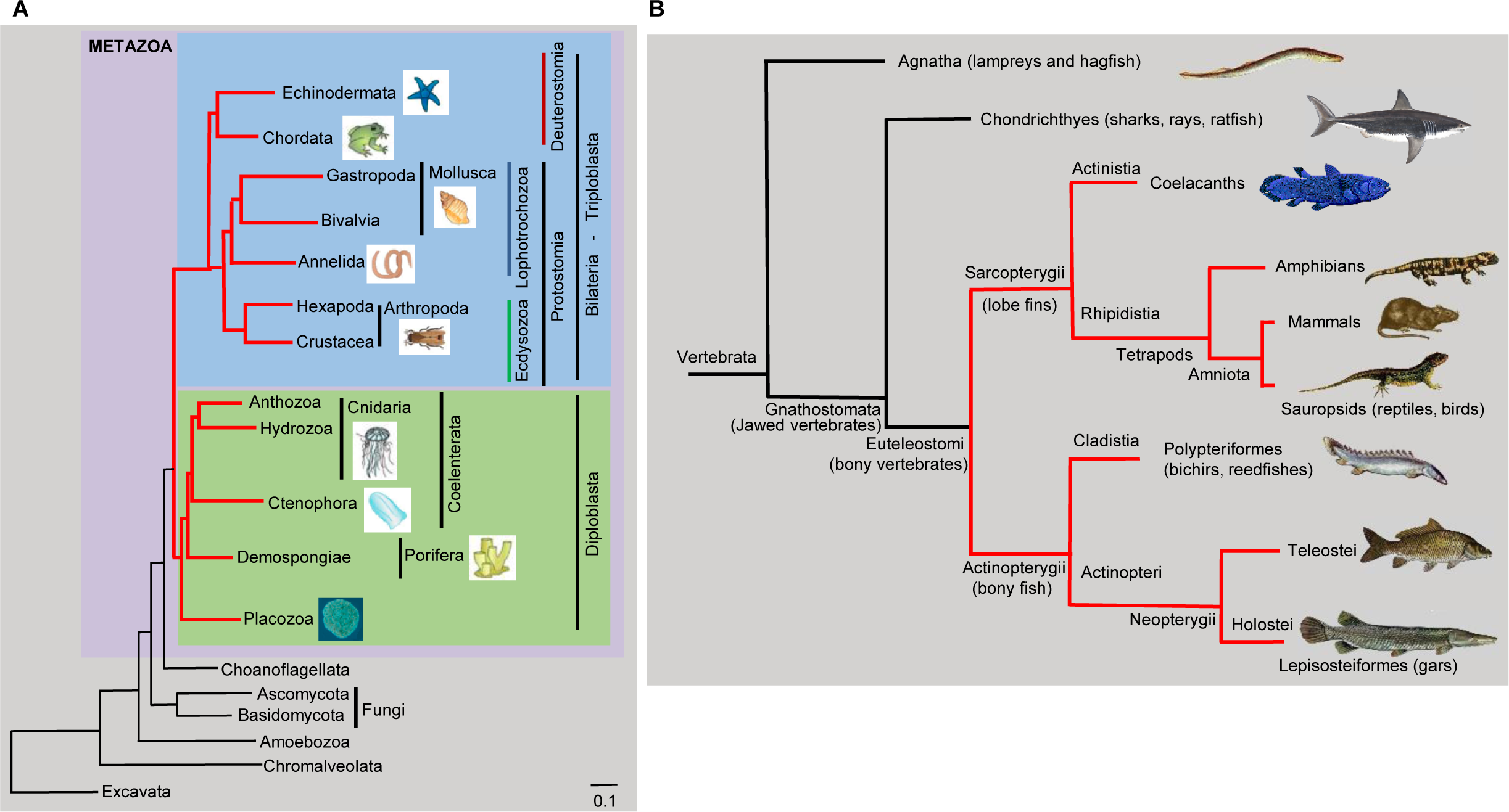
Metazoan origin of *NRBP* and the emergence of the paralog *NRBP2* in the Euteleostomi, related to Figure 7. **(A)** We mapped 2,065 homologs of human *NRBP1/2* to a phylogenetic tree of eukaryotes. Clades drawn with red lines indicate the presence of *NRBP*, whereas clades drawn with black lines indicate the absence of *NRBP*. *NRBP* is present in the early metazoan branches Porifera and Placozoa, but it is absent from all non-metazoan eukaryotes. The figure is a modified image from ^88^. The original image is licensed under the Creative Commons Attribution 2.5 Generic license (https://creativecommons.org/licenses/by/2.5/deed.en). **(B)** We mapped 669 candidate orthologues of human *NRBP2* to a phylogenetic tree of vertebrates.^89^ Clades drawn with red lines indicate the presence of *NRBP2*, whereas clades drawn with black lines indicate the absence of *NRBP2*. All *NRBP2* map either to the Sarcopterygii or to the Actinopterygii, indicating an evolutionary origin of *NRBP2* either in the Eutelestomi, or earlier in vertebrate evolution. All presented clades (black and red lines) have candidate orthologues of *NRBP1*. Remarkably, clades with *NRBP2* have a ‘Type II’ L1 ORF1, which is characterised by ‘transposase 22’, whereas the clades lacking *NRBP2* also lack the transposase 22. For details, please refer to the discussion section. The image is modified from https://en.wikipedia.org/wiki/Actinopterygii.

## Supplemental Tables

**Table S1. Analysis of NRBP1/2 interactors, related to Figure 1**. The list includes interactors of NRBP1/2 identified in MS, fold change ≥ 5, p < 0.05. Overlap of the NRBP1/2 interactors with known ORF1 interactors and proteins that regulate L1. GO term analysis of NRBP1/2 interactors using DAVID.

**Table S2. Analysis of RNAseq data upon knockdown of NRBP1 or NRBP2, related to Figure 6**. GO terms analysis was performed by using DAVID.

**Table S3. Sequences, antibodies and key solutions.**

## References

1. Otto, S.P., and Yong, P. (2002). The evolution of gene duplicates. Adv Genet 46, 451–483. 10.1016/s0065-2660(02)46017-8.

2. Innan, H., and Kondrashov, F. (2010). The evolution of gene duplications: classifying and distinguishing between models. Nat Rev Genet 11, 97–108. 10.1038/nrg2689.

3. Zhang, Y., Duc, A.C., Rao, S., Sun, X.L., Bilbee, A.N., Rhodes, M., Li, Q., Kappes, D.J., Rhodes, J., and Wiest, D.L. (2013). Control of hematopoietic stem cell emergence by antagonistic functions of ribosomal protein paralogs. Dev Cell 24, 411–425. 10.1016/j.devcel.2013.01.018.

4. Shum, E.Y., Jones, S.H., Shao, A., Chousal, J.N., Krause, M.D., Chan, W.K., Lou, C.H., Espinoza, J.L., Song, H.W., Phan, M.H., et al. (2016). The Antagonistic Gene Paralogs Upf3a and Upf3b Govern Nonsense-Mediated RNA Decay. Cell 165, 382–395. 10.1016/j.cell.2016.02.046.

5. Reddy, K.C., Dror, T., Underwood, R.S., Osman, G.A., Elder, C.R., Desjardins, C.A., Cuomo, C.A., Barkoulas, M., and Troemel, E.R. (2019). Antagonistic paralogs control a switch between growth and pathogen resistance in C. elegans. PLoS Pathog 15, e1007528. 10.1371/journal.ppat.1007528.

6. Riegel, K., Vijayarangakannan, P., Kechagioglou, P., Bogucka, K., and Rajalingam, K. (2022). Recent advances in targeting protein kinases and pseudokinases in cancer biology. Front Cell Dev Biol 10, 942500. 10.3389/fcell.2022.942500.

7. Kerr, J.S., and Wilson, C.H. (2013). Nuclear receptor-binding protein 1: a novel tumour suppressor and pseudokinase. Biochem Soc Trans 41, 1055–1060. 10.1042/BST20130069.

8. Lim, R., Winteringham, L.N., Williams, J.H., McCulloch, R.K., Ingley, E., Tiao, J.Y., Lalonde, J.P., Tsai, S., Tilbrook, P.A., Sun, Y., et al. (2002). MADM, a novel adaptor protein that mediates phosphorylation of the 14-3-3 binding site of myeloid leukemia factor 1. J Biol Chem 277, 40997–41008. 10.1074/jbc.M206041200.

9. Hooper, J.D., Baker, E., Ogbourne, S.M., Sutherland, G.R., and Antalis, T.M. (2000). Cloning of the cDNA and localization of the gene encoding human NRBP, a ubiquitously expressed, multidomain putative adapter protein. Genomics 66, 113–118. 10.1006/geno.2000.6167.

10. Wilson, C.H., Crombie, C., van der Weyden, L., Poulogiannis, G., Rust, A.G., Pardo, M., Gracia, T., Yu, L., Choudhary, J., Poulin, G.B., et al. (2012). Nuclear receptor binding protein 1 regulates intestinal progenitor cell homeostasis and tumour formation. EMBO J 31, 2486–2497. 10.1038/emboj.2012.91.

11. Yasukawa, T., Tsutsui, A., Tomomori-Sato, C., Sato, S., Saraf, A., Washburn, M.P., Florens, L., Terada, T., Shimizu, K., Conaway, R.C., et al. (2020). NRBP1-Containing CRL2/CRL4A Regulates Amyloid beta Production by Targeting BRI2 and BRI3 for Degradation. Cell Rep 30, 3478–3491 e3476. 10.1016/j.celrep.2020.02.059.

12. Yang, X., Cruz, M.I., Nguyen, E.V., Huang, C., Schittenhelm, R.B., Luu, J., Cowley, K.J., Shin, S.Y., Nguyen, L.K., Lim Kam Sian, T.C.C., et al. (2023). The pseudokinase NRBP1 activates Rac1/Cdc42 via P-Rex1 to drive oncogenic signalling in triple-negative breast cancer. Oncogene 42, 833–847. 10.1038/s41388-023-02594-w.

13. Hwang, J., Haque, M.A., Suzuki, H., Dijke, P.T., and Kato, M. (2020). THG-1 suppresses SALL4 degradation to induce stemness genes and tumorsphere formation through antagonizing NRBP1 in squamous cell carcinoma cells. Biochem Biophys Res Commun 523, 307–314. 10.1016/j.bbrc.2019.11.149.

14. Larsson, J., Forsberg, M., Brannvall, K., Zhang, X.Q., Enarsson, M., Hedborg, F., and Forsberg-Nilsson, K. (2008). Nuclear receptor binding protein 2 is induced during neural progenitor differentiation and affects cell survival. Mol Cell Neurosci 39, 32–39. 10.1016/j.mcn.2008.05.013.

15. Zhang, L., Ge, C., Zhao, F., Zhang, Y., Wang, X., Yao, M., and Li, J. (2016). NRBP2 Overexpression Increases the Chemosensitivity of Hepatocellular Carcinoma Cells via Akt Signaling. Cancer Res 76, 7059–7071. 10.1158/0008-5472.CAN-16-0937.

16. Li, Z., Liu, B., Li, C., Sun, S., Zhang, H., Sun, S., Wang, Z., and Zhang, X. (2021). NRBP2 Functions as a Tumor Suppressor and Inhibits Epithelial-to-Mesenchymal Transition in Breast Cancer. Front Oncol 11, 634026. 10.3389/fonc.2021.634026.

17. Lander, E.S., Linton, L.M., Birren, B., Nusbaum, C., Zody, M.C., Baldwin, J., Devon, K., Dewar, K., Doyle, M., FitzHugh, W., et al. (2001). Initial sequencing and analysis of the human genome. Nature 409, 860–921. 10.1038/35057062.

18. Beck, C.R., Garcia-Perez, J.L., Badge, R.M., and Moran, J.V. (2011). LINE-1 elements in structural variation and disease. Annu Rev Genomics Hum Genet 12, 187–215. 10.1146/annurev-genom-082509-141802.

19. Gorbunova, V., Seluanov, A., Mita, P., McKerrow, W., Fenyo, D., Boeke, J.D., Linker, S.B., Gage, F.H., Kreiling, J.A., Petrashen, A.P., et al. (2021). The role of retrotransposable elements in ageing and age-associated diseases. Nature 596, 43–53. 10.1038/s41586-021-03542-y.

20. Mathavarajah, S., and Dellaire, G. (2024). LINE-1: an emerging initiator of cGAS-STING signalling and inflammation that is dysregulated in disease. Biochem Cell Biol 102, 38–46. 10.1139/bcb-2023-0134.

21. Feng, Q., Moran, J.V., Kazazian, H.H., Jr., and Boeke, J.D. (1996). Human L1 retrotransposon encodes a conserved endonuclease required for retrotransposition. Cell 87, 905–916. 10.1016/s0092-8674(00)81997-2.

22. Holmes, S.E., Singer, M.F., and Swergold, G.D. (1992). Studies on p40, the leucine zipper motif-containing protein encoded by the first open reading frame of an active human LINE-1 transposable element. J Biol Chem 267, 19765–19768.

23. Moran, J.V., Holmes, S.E., Naas, T.P., DeBerardinis, R.J., Boeke, J.D., and Kazazian, H.H., Jr. (1996). High frequency retrotransposition in cultured mammalian cells. Cell 87, 917–927. 10.1016/s0092-8674(00)81998-4.

24. Ergun, S., Buschmann, C., Heukeshoven, J., Dammann, K., Schnieders, F., Lauke, H., Chalajour, F., Kilic, N., Stratling, W.H., and Schumann, G.G. (2004). Cell type-specific expression of LINE-1 open reading frames 1 and 2 in fetal and adult human tissues. J Biol Chem 279, 27753–27763. 10.1074/jbc.M312985200.

25. Mathias, S.L., Scott, A.F., Kazazian, H.H., Jr., Boeke, J.D., and Gabriel, A. (1991). Reverse transcriptase encoded by a human transposable element. Science 254, 1808–1810. 10.1126/science.1722352.

26. Hohjoh, H., and Singer, M.F. (1996). Cytoplasmic ribonucleoprotein complexes containing human LINE-1 protein and RNA. EMBO J 15, 630–639.

27. Kulpa, D.A., and Moran, J.V. (2005). Ribonucleoprotein particle formation is necessary but not sufficient for LINE-1 retrotransposition. Hum Mol Genet 14, 3237–3248. 10.1093/hmg/ddi354.

28. Kulpa, D.A., and Moran, J.V. (2006). Cis-preferential LINE-1 reverse transcriptase activity in ribonucleoprotein particles. Nat Struct Mol Biol 13, 655–660. 10.1038/nsmb1107.

29. Cost, G.J., Feng, Q., Jacquier, A., and Boeke, J.D. (2002). Human L1 element target-primed reverse transcription in vitro. EMBO J 21, 5899–5910. 10.1093/emboj/cdf592.

30. Ariumi, Y. (2016). Guardian of the Human Genome: Host Defense Mechanisms against LINE-1 Retrotransposition. Frontiers in chemistry 4, 28. 10.3389/fchem.2016.00028.

31. Goodier, J.L., Cheung, L.E., and Kazazian, H.H., Jr. (2013). Mapping the LINE1 ORF1 protein interactome reveals associated inhibitors of human retrotransposition. Nucleic Acids Res 41, 7401–7419. 10.1093/nar/gkt512.

32. Taylor, M.S., LaCava, J., Mita, P., Molloy, K.R., Huang, C.R., Li, D., Adney, E.M., Jiang, H., Burns, K.H., Chait, B.T., et al. (2013). Affinity proteomics reveals human host factors implicated in discrete stages of LINE-1 retrotransposition. Cell 155, 1034–1048. 10.1016/j.cell.2013.10.021.

33. Percharde, M., Lin, C.J., Yin, Y., Guan, J., Peixoto, G.A., Bulut-Karslioglu, A., Biechele, S., Huang, B., Shen, X., and Ramalho-Santos, M. (2018). A LINE1-Nucleolin Partnership Regulates Early Development and ESC Identity. Cell 174, 391–405 e319. 10.1016/j.cell.2018.05.043.

34. Richardson, S.R., Gerdes, P., Gerhardt, D.J., Sanchez-Luque, F.J., Bodea, G.O., Munoz-Lopez, M., Jesuadian, J.S., Kempen, M.H.C., Carreira, P.E., Jeddeloh, J.A., et al. (2017). Heritable L1 retrotransposition in the mouse primordial germline and early embryo. Genome Res 27, 1395–1405. 10.1101/gr.219022.116.

35. Seleme, M.C., Vetter, M.R., Cordaux, R., Bastone, L., Batzer, M.A., and Kazazian, H.H., Jr. (2006). Extensive individual variation in L1 retrotransposition capability contributes to human genetic diversity. Proc Natl Acad Sci U S A 103, 6611–6616. 10.1073/pnas.0601324103.

36. Mangoni, D., Simi, A., Lau, P., Armaos, A., Ansaloni, F., Codino, A., Damiani, D., Floreani, L., Di Carlo, V., Vozzi, D., et al. (2023). LINE-1 regulates cortical development by acting as long non-coding RNAs. Nat Commun 14, 4974. 10.1038/s41467-023-40743-7.

37. Taylor, M.S., Altukhov, I., Molloy, K.R., Mita, P., Jiang, H., Adney, E.M., Wudzinska, A., Badri, S., Ischenko, D., Eng, G., et al. (2018). Dissection of affinity captured LINE-1 macromolecular complexes. Elife 7. 10.7554/eLife.30094.

38. Luqman-Fatah, A., Watanabe, Y., Uno, K., Ishikawa, F., Moran, J.V., and Miyoshi, T. (2023). The interferon stimulated gene-encoded protein HELZ2 inhibits human LINE-1 retrotransposition and LINE-1 RNA-mediated type I interferon induction. Nat Commun 14, 203. 10.1038/s41467-022-35757-6.

39. Esnault, C., Heidmann, O., Delebecque, F., Dewannieux, M., Ribet, D., Hance, A.J., Heidmann, T., and Schwartz, O. (2005). APOBEC3G cytidine deaminase inhibits retrotransposition of endogenous retroviruses. Nature 433, 430–433. 10.1038/nature03238.

40. Hu, S., Li, J., Xu, F., Mei, S., Le Duff, Y., Yin, L., Pang, X., Cen, S., Jin, Q., Liang, C., and Guo, F. (2015). SAMHD1 Inhibits LINE-1 Retrotransposition by Promoting Stress Granule Formation. PLoS Genet 11, e1005367. 10.1371/journal.pgen.1005367.

41. Dai, L., Taylor, M.S., O’Donnell, K.A., and Boeke, J.D. (2012). Poly(A) binding protein C1 is essential for efficient L1 retrotransposition and affects L1 RNP formation. Mol Cell Biol 32, 4323–4336. 10.1128/MCB.06785-11.

42. De Langhe, S., Haataja, L., Senadheera, D., Groffen, J., and Heisterkamp, N. (2002). Interaction of the small GTPase Rac3 with NRBP, a protein with a kinase-homology domain. Int J Mol Med 9, 451–459.

43. Wootton, J.C., and Federhen, S. (1996). Analysis of compositionally biased regions in sequence databases. Methods Enzymol 266, 554–571. 10.1016/s0076-6879(96)66035-2.

44. Jumper, J., Evans, R., Pritzel, A., Green, T., Figurnov, M., Ronneberger, O., Tunyasuvunakool, K., Bates, R., Zidek, A., Potapenko, A., et al. (2021). Highly accurate protein structure prediction with AlphaFold. Nature 596, 583–589. 10.1038/s41586-021-03819-2.

45. Varadi, M., Bertoni, D., Magana, P., Paramval, U., Pidruchna, I., Radhakrishnan, M., Tsenkov, M., Nair, S., Mirdita, M., Yeo, J., et al. (2024). AlphaFold Protein Structure Database in 2024: providing structure coverage for over 214 million protein sequences. Nucleic Acids Res 52, D368–D375. 10.1093/nar/gkad1011.

46. De Cecco, M., Ito, T., Petrashen, A.P., Elias, A.E., Skvir, N.J., Criscione, S.W., Caligiana, A., Brocculi, G., Adney, E.M., Boeke, J.D., et al. (2019). L1 drives IFN in senescent cells and promotes age-associated inflammation. Nature 566, 73–78. 10.1038/s41586-018-0784-9.

47. Lagisquet, J., Zuber, K., and Gramberg, T. (2021). Recognize Yourself-Innate Sensing of Non-LTR Retrotransposons. Viruses 13. 10.3390/v13010094.

48. Zhao, K., Du, J., Peng, Y., Li, P., Wang, S., Wang, Y., Hou, J., Kang, J., Zheng, W., Hua, S., and Yu, X.F. (2018). LINE1 contributes to autoimmunity through both RIG-I- and MDA5-mediated RNA sensing pathways. J Autoimmun 90, 105–115. 10.1016/j.jaut.2018.02.007.

49. Tunbak, H., Enriquez-Gasca, R., Tie, C.H.C., Gould, P.A., Mlcochova, P., Gupta, R.K., Fernandes, L., Holt, J., van der Veen, A.G., Giampazolias, E., et al. (2020). The HUSH complex is a gatekeeper of type I interferon through epigenetic regulation of LINE-1s. Nat Commun 11, 5387. 10.1038/s41467-020-19170-5.

50. Thomas, C.A., Tejwani, L., Trujillo, C.A., Negraes, P.D., Herai, R.H., Mesci, P., Macia, A., Crow, Y.J., and Muotri, A.R. (2017). Modeling of TREX1-Dependent Autoimmune Disease using Human Stem Cells Highlights L1 Accumulation as a Source of Neuroinflammation. Cell Stem Cell 21, 319–331 e318. 10.1016/j.stem.2017.07.009.

51. Zhao, K., Du, J., Han, X., Goodier, J.L., Li, P., Zhou, X., Wei, W., Evans, S.L., Li, L., Zhang, W., et al. (2013). Modulation of LINE-1 and Alu/SVA retrotransposition by Aicardi-Goutieres syndrome-related SAMHD1. Cell Rep 4, 1108–1115. 10.1016/j.celrep.2013.08.019.

52. Orecchini, E., Doria, M., Antonioni, A., Galardi, S., Ciafre, S.A., Frassinelli, L., Mancone, C., Montaldo, C., Tripodi, M., and Michienzi, A. (2017). ADAR1 restricts LINE-1 retrotransposition. Nucleic Acids Res 45, 155–168. 10.1093/nar/gkw834.

53. Guo, Y., Walsh, A.M., Fearon, U., Smith, M.D., Wechalekar, M.D., Yin, X., Cole, S., Orr, C., McGarry, T., Canavan, M., et al. (2017). CD40L-Dependent Pathway Is Active at Various Stages of Rheumatoid Arthritis Disease Progression. J Immunol 198, 4490–4501. 10.4049/jimmunol.1601988.

54. Walsh, A.M., Wechalekar, M.D., Guo, Y., Yin, X., Weedon, H., Proudman, S.M., Smith, M.D., and Nagpal, S. (2017). Triple DMARD treatment in early rheumatoid arthritis modulates synovial T cell activation and plasmablast/plasma cell differentiation pathways. PLoS One 12, e0183928. 10.1371/journal.pone.0183928.

55. Kwon, A., Scott, S., Taujale, R., Yeung, W., Kochut, K.J., Eyers, P.A., and Kannan, N. (2019). Tracing the origin and evolution of pseudokinases across the tree of life. Sci Signal 12. 10.1126/scisignal.aav3810.

56. Qi, W., Yan, Y., Pfeifer, D., Donner, V.G.E., Wang, Y., Maier, W., and Baumeister, R. (2017). C. elegans DAF-16/FOXO interacts with TGF-ss/BMP signaling to induce germline tumor formation via mTORC1 activation. PLoS Genet 13, e1006801. 10.1371/journal.pgen.1006801.

57. Ivancevic, A.M., Kortschak, R.D., Bertozzi, T., and Adelson, D.L. (2016). LINEs between Species: Evolutionary Dynamics of LINE-1 Retrotransposons across the Eukaryotic Tree of Life. Genome Biol Evol 8, 3301–3322. 10.1093/gbe/evw243.

58. Mistry, J., Chuguransky, S., Williams, L., Qureshi, M., Salazar, G.A., Sonnhammer, E.L.L., Tosatto, S.C.E., Paladin, L., Raj, S., Richardson, L.J., et al. (2021). Pfam: The protein families database in 2021. Nucleic Acids Res 49, D412–D419. 10.1093/nar/gkaa913.

59. Liu, Q., Yi, D., Ding, J., Mao, Y., Wang, S., Ma, L., Li, Q., Wang, J., Zhang, Y., Zhao, J., et al. (2023). MOV10 recruits DCP2 to decap human LINE-1 RNA by forming large cytoplasmic granules with phase separation properties. EMBO Rep 24, e56512. 10.15252/embr.202256512.

60. Thedieck, K., Polak, P., Kim, M.L., Molle, K.D., Cohen, A., Jeno, P., Arrieumerlou, C., and Hall, M.N. (2007). PRAS40 and PRR5-like protein are new mTOR interactors that regulate apoptosis. PLoS One 2, e1217. 10.1371/journal.pone.0001217.

61. Schwarz, J.J., Wiese, H., Tolle, R.C., Zarei, M., Dengjel, J., Warscheid, B., and Thedieck, K. (2015). Functional Proteomics Identifies Acinus L as a Direct Insulin- and Amino Acid-Dependent Mammalian Target of Rapamycin Complex 1 (mTORC1) Substrate. Mol Cell Proteomics 14, 2042–2055. 10.1074/mcp.M114.045807.

62. Cox, J., and Mann, M. (2008). MaxQuant enables high peptide identification rates, individualized p.p.b.-range mass accuracies and proteome-wide protein quantification. Nat Biotechnol 26, 1367–1372. 10.1038/nbt.1511.

63. Tyanova, S., Temu, T., and Cox, J. (2016). The MaxQuant computational platform for mass spectrometry-based shotgun proteomics. Nat Protoc 11, 2301–2319. 10.1038/nprot.2016.136.

64. Cox, J., Hein, M.Y., Luber, C.A., Paron, I., Nagaraj, N., and Mann, M. (2014). Accurate proteome-wide label-free quantification by delayed normalization and maximal peptide ratio extraction, termed MaxLFQ. Mol Cell Proteomics 13, 2513–2526. 10.1074/mcp.M113.031591.

65. Tyanova, S., Temu, T., Sinitcyn, P., Carlson, A., Hein, M.Y., Geiger, T., Mann, M., and Cox, J. (2016). The Perseus computational platform for comprehensive analysis of (prote)omics data. Nat Methods 13, 731–740. 10.1038/nmeth.3901.

66. Perez-Riverol, Y., Csordas, A., Bai, J., Bernal-Llinares, M., Hewapathirana, S., Kundu, D.J., Inuganti, A., Griss, J., Mayer, G., Eisenacher, M., et al. (2019). The PRIDE database and related tools and resources in 2019: improving support for quantification data. Nucleic Acids Res 47, D442–D450. 10.1093/nar/gky1106.

67. Dewannieux, M., Dupressoir, A., Harper, F., Pierron, G., and Heidmann, T. (2004). Identification of autonomous IAP LTR retrotransposons mobile in mammalian cells. Nat Genet 36, 534–539. 10.1038/ng1353.

68. Livak, K.J., and Schmittgen, T.D. (2001). Analysis of relative gene expression data using real-time quantitative PCR and the 2(-Delta Delta C(T)) Method. Methods 25, 402–408. 10.1006/meth.2001.1262.

69. Martin, M. (2011). Cutadapt removes adapter sequences from high-throughput sequencing reads. EMBnet.journal 17. DOI: 10.14806/ej.17.1.200.

70. Dobin, A., Davis, C.A., Schlesinger, F., Drenkow, J., Zaleski, C., Jha, S., Batut, P., Chaisson, M., and Gingeras, T.R. (2013). STAR: ultrafast universal RNA-seq aligner. Bioinformatics 29, 15–21. 10.1093/bioinformatics/bts635.

71. Liao, Y., Smyth, G.K., and Shi, W. (2014). featureCounts: an efficient general purpose program for assigning sequence reads to genomic features. Bioinformatics 30, 923–930. 10.1093/bioinformatics/btt656.

72. Robinson, M.D., McCarthy, D.J., and Smyth, G.K. (2010). edgeR: a Bioconductor package for differential expression analysis of digital gene expression data. Bioinformatics 26, 139–140. 10.1093/bioinformatics/btp616.

73. Guo, H., Chitiprolu, M., Gagnon, D., Meng, L., Perez-Iratxeta, C., Lagace, D., and Gibbings, D. (2014). Autophagy supports genomic stability by degrading retrotransposon RNA. Nat Commun 5, 5276. 10.1038/ncomms6276.

74. UniProt, C. (2023). UniProt: the Universal Protein Knowledgebase in 2023. Nucleic Acids Res 51, D523–D531. 10.1093/nar/gkac1052.

75. Camacho, C., Coulouris, G., Avagyan, V., Ma, N., Papadopoulos, J., Bealer, K., and Madden, T.L. (2009). BLAST+: architecture and applications. BMC Bioinformatics 10, 421. 10.1186/1471-2105-10-421.

76. Boutet, E., Lieberherr, D., Tognolli, M., Schneider, M., and Bairoch, A. (2007). UniProtKB/Swiss-Prot. Methods Mol Biol 406, 89–112. 10.1007/978-1-59745-535-0_4.

77. Leinonen, R., Akhtar, R., Birney, E., Bower, L., Cerdeno-Tarraga, A., Cheng, Y., Cleland, I., Faruque, N., Goodgame, N., Gibson, R., et al. (2011). The European Nucleotide Archive. Nucleic Acids Res 39, D28–31. 10.1093/nar/gkq967.

78. Katoh, K., and Standley, D.M. (2013). MAFFT multiple sequence alignment software version 7: improvements in performance and usability. Mol Biol Evol 30, 772–780. 10.1093/molbev/mst010.

79. Capella-Gutierrez, S., Silla-Martinez, J.M., and Gabaldon, T. (2009). trimAl: a tool for automated alignment trimming in large-scale phylogenetic analyses. Bioinformatics 25, 1972–1973. 10.1093/bioinformatics/btp348.

80. Minh, B.Q., Schmidt, H.A., Chernomor, O., Schrempf, D., Woodhams, M.D., von Haeseler, A., and Lanfear, R. (2020). IQ-TREE 2: New Models and Efficient Methods for Phylogenetic Inference in the Genomic Era. Mol Biol Evol 37, 1530–1534. 10.1093/molbev/msaa015.

81. Hoang, D.T., Chernomor, O., von Haeseler, A., Minh, B.Q., and Vinh, L.S. (2018). UFBoot2: Improving the Ultrafast Bootstrap Approximation. Mol Biol Evol 35, 518–522. 10.1093/molbev/msx281.

82. Kalyaanamoorthy, S., Minh, B.Q., Wong, T.K.F., von Haeseler, A., and Jermiin, L.S. (2017). ModelFinder: fast model selection for accurate phylogenetic estimates. Nat Methods 14, 587–589. 10.1038/nmeth.4285.

83. Letunic, I., and Bork, P. (2024). Interactive Tree of Life (iTOL) v6: recent updates to the phylogenetic tree display and annotation tool. Nucleic Acids Res. 10.1093/nar/gkae268.

84. Talevich, E., Invergo, B.M., Cock, P.J., and Chapman, B.A. (2012). Bio.Phylo: a unified toolkit for processing, analyzing and visualizing phylogenetic trees in Biopython. BMC Bioinformatics 13, 209. 10.1186/1471-2105-13-209.

85. Huntley, M.A., and Golding, G.B. (2002). Simple sequences are rare in the Protein Data Bank. Proteins 48, 134–140. 10.1002/prot.10150.

86. Wang, X., Wang, X., Zhang, H., Lv, M., Zuo, T., Wu, H., Wang, J., Liu, D., Wang, C., Zhang, J., et al. (2013). Interactions between HIV-1 Vif and human ElonginB-ElonginC are important for CBF-beta binding to Vif. Retrovirology 10, 94. 10.1186/1742-4690-10-94.

87. Schieven, S.M., Traets, J.J.H., Vliet, A.V., Baalen, M.V., Song, J.Y., Guimaraes, M.D.S., Kuilman, T., and Peeper, D.S. (2023). The Elongin BC Complex Negatively Regulates AXL and Marks a Differentiated Phenotype in Melanoma. Mol Cancer Res 21, 428–443. 10.1158/1541-7786.MCR-22-0648.

88. Schierwater, B., Eitel, M., Jakob, W., Osigus, H.J., Hadrys, H., Dellaporta, S.L., Kolokotronis, S.O., and Desalle, R. (2009). Concatenated analysis sheds light on early metazoan evolution and fuels a modern “urmetazoon” hypothesis. PLoS Biol 7, e20. 10.1371/journal.pbio.1000020.

89. Betancur, R.R., Broughton, R.E., Wiley, E.O., Carpenter, K., Lopez, J.A., Li, C., Holcroft, N.I., Arcila, D., Sanciangco, M., Cureton Ii, J.C., et al. (2013). The tree of life and a new classification of bony fishes. PLoS Curr 5. 10.1371/currents.tol.53ba26640df0ccaee75bb165c8c26288.

